# The *homie* insulator has sub-elements with different insulating and long-range pairing properties

**DOI:** 10.1101/2024.02.01.578481

**Authors:** Miki Fujioka, Wenfan Ke, Paul Schedl, James B Jaynes

## Abstract

Chromatin insulators are major determinants of chromosome architecture. Specific architectures induced by insulators profoundly influence nuclear processes, including how enhancers and promoters interact over long distances and between homologous chromosomes. Insulators can pair with copies of themselves in *trans* to facilitate homolog pairing. They can also pair with other insulators, sometimes with great specificity, inducing long-range chromosomal loops. Contrary to their canonical function of enhancer blocking, these loops can bring distant enhancers and promoters together to activate gene expression, while at the same time blocking other interactions in *cis*. The details of these effects depend on the choice of pairing partner, and on the orientation specificity of pairing, implicating the 3-dimensional architecture as a major functional determinant. Here we dissect the *homie* insulator from the Drosophila *even skipped* (*eve*) locus, to understand its substructure. We test pairing function based on *homie*-carrying transgenes interacting with endogenous *eve*. The assay is sensitive to both pairing strength and orientation. Using this assay, we found that a Su(Hw) binding site in *homie* is required for efficient long-range interaction, although some activity remains without it. This binding site also contributes to the canonical insulator activities of enhancer blocking and barrier function. Based on this and other results from our functional dissection, each of the canonical insulator activities, chromosomal loop formation, enhancer blocking, and barrier activity, are partially separable. Our results show the complexity inherent in insulator functions, which can be provided by an array of different proteins with both shared and distinct properties.

**GRAPHICAL ABSTRACT:** 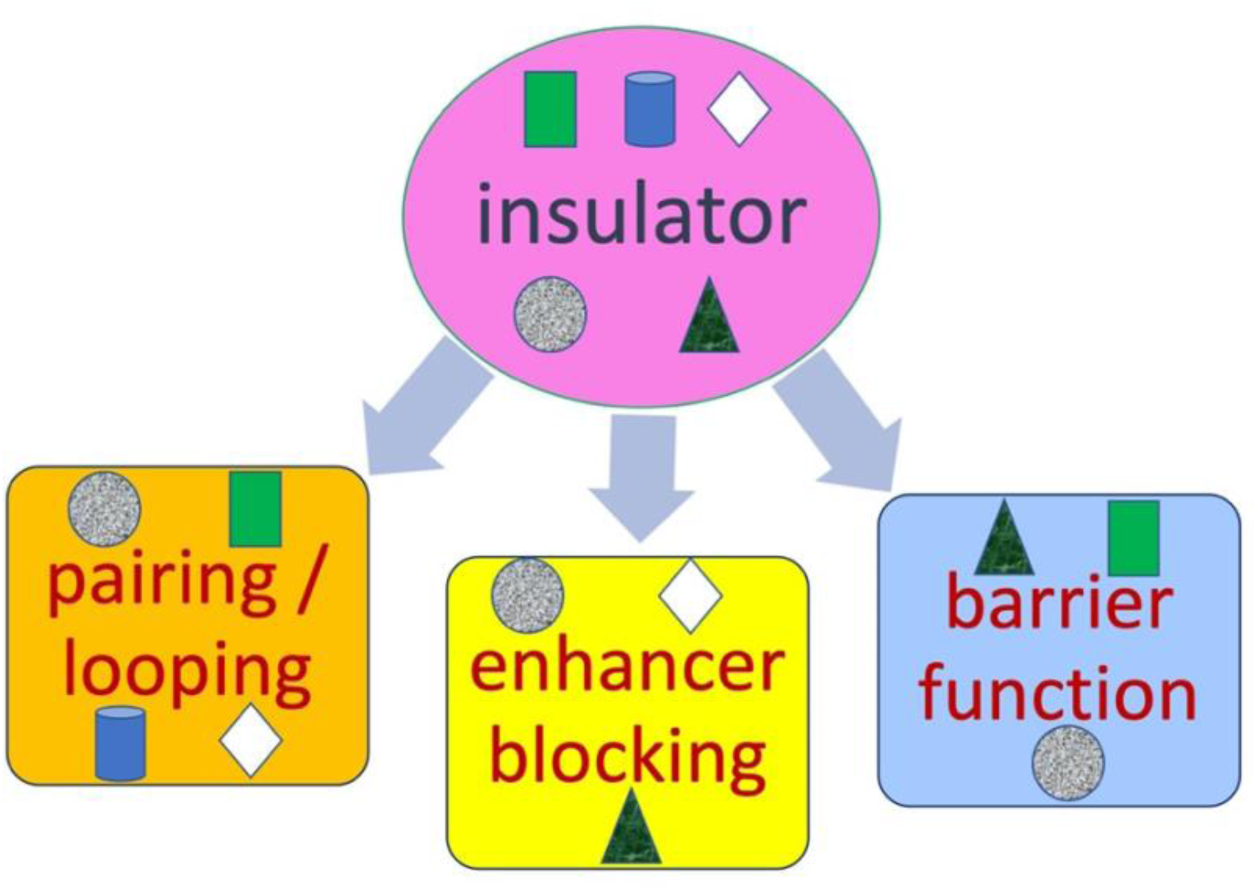

## Introduction

Genomic elements called boundaries or insulators function to separate genes from each other, or, in some cases, to separate distinct regulatory domains within a gene (e.g., in the Hox gene complexes).

However, they have properties that have proven to be difficult to reconcile mechanistically. For example, while these elements can block enhancer-promoter interactions when inserted between regulatory elements, they can also mediate long-distance regulatory interactions (Chetverina et al. 2017; Özdemir and Gambetta 2019; Schwartz and Cavalli 2017).

One of the first insulator elements discovered is located within the *Drosophila gypsy* retrotransposon (Bender et al. 1983; Modolell et al. 1983). The *gypsy* transposon was found to be responsible for many spontaneous mutations in flies. In many cases, the mutant phenotypes arose because the *gypsy* transposon inserted between a gene and its regulatory elements, blocking regulatory interactions (Geyer et al. 1986; Jack et al. 1991; Parkhurst and Corces 1985; Peifer and Bender 1986). The insulator properties of *gypsy* have been attributed to the binding of a set of chromatin proteins, the sequence-specific binding component of which is Su(Hw) (Suppressor of Hairy Wing) (Geyer and Corces 1992; Holdridge and Dorsett 1991; Mallin et al. 1998; Parkhurst et al. 1988; Roseman et al. 1993; Spana et al. 1988). Genetic and biochemical analysis identified Mod(mdg4) (Gause et al. 2001; Gerasimova et al. 1995; Ghosh et al. 2001) and Cp190 (Pai et al. 2004) as Su(Hw)-interacting proteins. While Su(Hw) is a zinc finger protein that binds DNA, both Mod(mdg4) and Cp190 contain BTB/POZ protein-protein interaction motifs. Studies have shown that the BTB/POZ domain of Cp190 interacts both with itself and with multiple other insulator binding proteins (Golovnin et al. 2023; Vogelmann et al. 2014), and Cp190 pull-down analysis identified additional insulator proteins in embryo nuclear extracts (Kaushal et al. 2022).

Several insulators found in the bithorax complex (BX-C) of Drosophila have been extensively studied, including *Mcp* (Karch et al. 1994), *Fab7* (Hagstrom et al. 1996; Karch et al. 1994), and *Fab8* (Barges et al. 2000; Zhou et al. 1999), as well as FS1 in the Antennapedia complex (Belozerov et al. 2003), and *scs* and *scs’*, which flank the *hsp70* locus (Kellum and Schedl 1991; Udvardy et al. 1985). Many insulator binding proteins that are known to contribute to insulator activity have been identified. These include Zw5 (Gaszner et al.

1999), GAF (Belozerov et al. 2003; Ohtsuki and Levine 1998), CTCF (Moon et al. 2005), BEAF-32 (Jiang et al. 2009; Roy et al. 2007; Zhao et al. 1995), Ibf1 and Ibf2 (Cuartero et al. 2014), Elba and Insensitive (Aoki et al. 2012; Fedotova et al. 2018), Pita and ZIPIC (Maksimenko et al. 2015; Zolotarev et al. 2016), M1BP (Bag et al. 2021; Li and Gilmour 2013), and Chromator (Gortchakov et al. 2005; Rath et al. 2004; Sexton et al. 2012).

Genome-wide analysis of the sequences associated with various insulator proteins suggested that there are different classes of insulators, based on binding by specific combinations of these proteins (Negre et al. 2010).

The Drosophila *even skipped* (*eve*) gene is flanked by two insulators, *nhomie* (*neighbor of homie*) and *homie* (*homing insulator at eve*), at its 5’- (upstream-) and 3’- (downstream-) ends, respectively (Figure 1a, b). These two elements define the *eve* TAD (topologically associating domain) (Bing et al. 2024; Fujioka et al. 2016; Fujioka et al. 2009; Ke et al. 2024). Genome-wide analysis showed that both *homie* and *nhomie* are bound by most of the insulator binding proteins mentioned above, as well as by Rad21, a component of the cohesin complex (http://chorogenome.ie-freiburg.mpg.de; Bag et al. 2021; Baxley et al. 2017; Cuartero et al. 2014; Cubeñas-Potts et al. 2016; Li and Gilmour 2013; Li et al. 2015; Maksimenko et al. 2015; Matzat et al. 2012; Ramírez et al. 2018; Schwartz et al. 2012; Soshnev et al. 2012; Van Bortle et al. 2014; Van Bortle et al. 2012; Wood et al. 2011; Zolotarev et al. 2016). The properties of *homie* have been well established. Like many other insulators, it has enhancer blocking activity (Fujioka et al. 2016; Fujioka et al. 2009) and an ability to prevent the spread of Polycomb-dependent silencing (Fujioka et al. 2013). *homie* abuts the promoter of an essential housekeeping gene, *TER94* (Leon and McKearin 1999; Pinter et al. 1998; Ruden et al. 2000).

**Figure 1.**
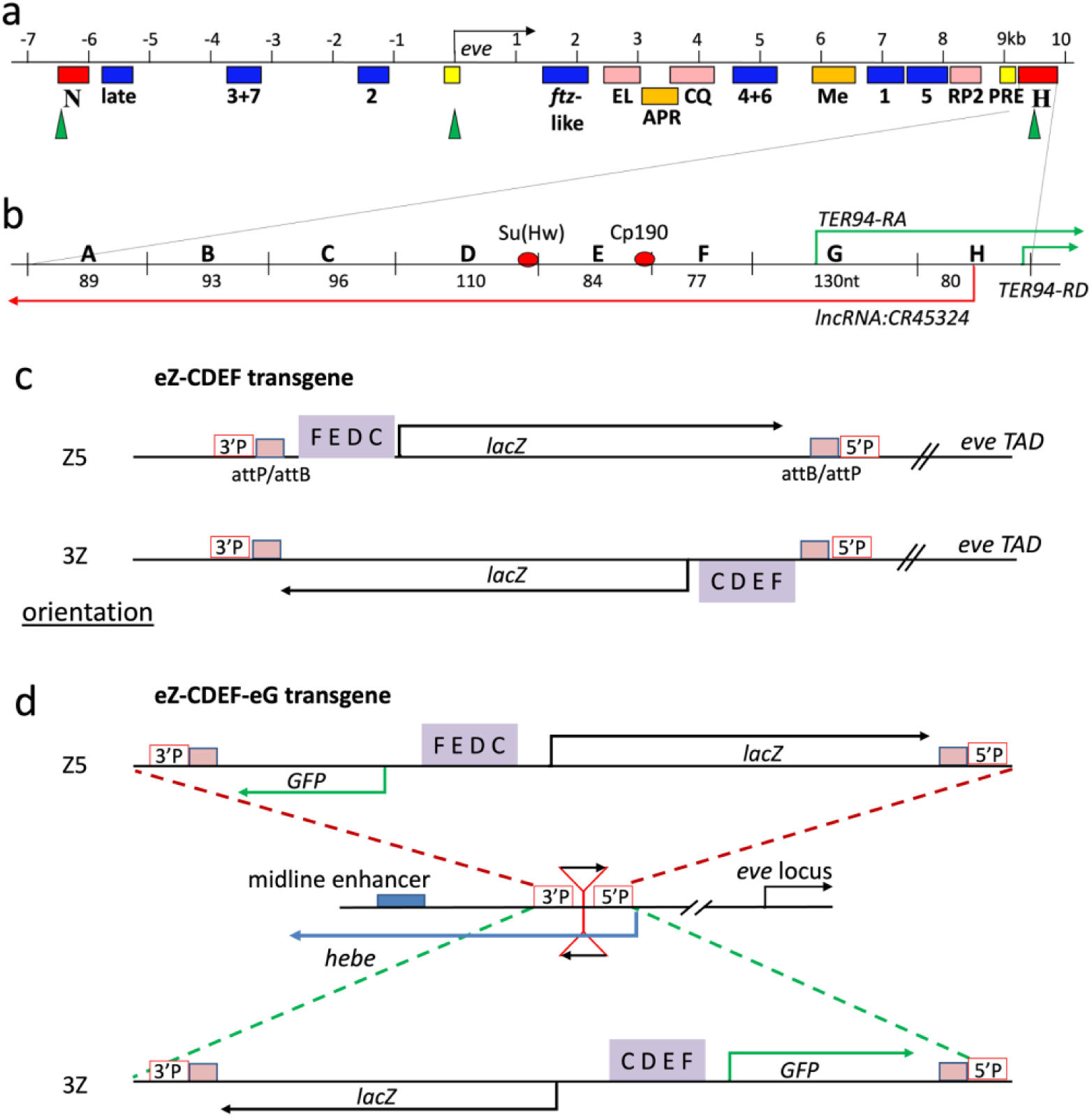
Maps of the *eve* TAD and transgenes inserted at –142kb from the *eve* locus. a: map of the *eve* locus. Blue: stripe enhancers (Fujioka et al. 1999; Sackerson et al. 1999; Small et al. 1992; Small et al. 1996) for the 7-“late”-stripe pattern (late), early stripes 3 and 7 (3+7), early stripe 2 (2), 7 *ftz*-like stripes (*ftz*-like), stripes 4 and 6 (4+6), stripe 1 (1), and stripe 5 (5). Pink: neuronal enhancers (Fujioka et al. 1999; Sackerson et al. 1999): EL (EL), CQ (CQ), RP2 + a/pCC (RP2) neurectodermal cells. Orange: tissue-specific enhancers (Fujioka et al. 1999; Sackerson et al. 1999): anal plate ring (APR), mesodermal cells (Me). Yellow: PRE (Fujioka et al. 2008). Red: insulators *nhomie* (N) and *homie* (H) (Fujioka et al. 2016; Fujioka et al. 2009). Green arrowheads: DNase I hypersensitive sites (Sackerson et al. 1999). **b: the *homie* sub-elements, ABCDEFGH.** The *TER94-RA* transcript starts in G. Both the *TER94-RD* transcript and the *lncRNA:CR45324* start in H (Flybase, Gramates et al. 2022). The positions of the Su(Hw) binding site and the Cp190-associated site (Baxley et al. 2017; Cuartero et al. 2014; Negre et al. 2010) are shown as red ovals. **c, d: the eZ-CDEF and eZ-CDEF-eG transgenes.** The –142kb attP site is located in the 5’ UTR of the *hebe* transcription unit. Because the attP site was originally inserted via P-element transposition, the 5’- and 3’-P-element ends flank it in the genome. The 3’-P-end (“3’P”) is located closer to the CNS midline enhancer of the *hebe* gene, while the 5’-P-end (“5’P”) is closer to the *eve* locus. The *hebe-RD* transcription unit (Flybase, Gramates et al. 2022) is shown as a blue arrow. Insertion of a transgene via RMCE results in two recombined versions of the original attP and attB sites (Bateman et al. 2006). Transgene insertion can occur in either orientation: “Z5”, where *lacZ* is closer to 5’P, or “3Z”, where *lacZ* is closer to 3’P. In the case of Z5, a modified *homie* is located between 3’P and the *eve-lacZ* reporter gene (*lacZ*: direction of transcription is shown as a black arrow). In the case of 3Z, modified *homie* is located between *eve-lacZ* and 5’P, as illustrated. The orientation of *homie* in the chromosome is indicated by the order of its sub-elements: either “CDEF” for the same orientation in the chromosome as endogenous *homie*, or “FEDC”, for the opposite orientation. Note that the two transgenes in C are identical except for their orientation in the chromosome, as are the two transgenes in D. The eZ- CDEF-eG vector is the same, except for the additional presence of an *eve-GFP* reporter (*GFP*, green arrow), transcribed in the opposite direction as *eve-lacZ*. Each reporter is driven by the *eve* basal promoter, which confers no *eve*-like expression on its own (Fujioka et al. 2016).

Since the *eve* TAD is assembled into a Polycomb-group (PcG)-silenced domain in most cells during all but the early stages of development (Negre et al. 2010; Schwartz et al. 2006; Tolhuis et al. 2006; Van Bortle et al.

2012), this barrier activity is thought to be a critically important function, as insulator protein binding sites are located at roughly half of the borders of Polycomb domains in *Drosophila* (De et al. 2020). In *eve-TER94* “pseudo-locus” transgenes, *homie* is required to keep the *eve* Polycomb response elements (PREs) from shutting down *TER94* through the spreading of the repressive PcG-dependent histone modification H3K27me3 (trimethylation of histone H3 residue lysine-27) (Fujioka et al. 2013).

In addition to these activities, *homie* and *nhomie* share another characteristic property of fly insulators, namely, an ability to physically pair with themselves and with each other (Bing et al. 2024; Chen et al. 2018; Fujioka et al. 2016; Fujioka et al. 2009; Ke et al. 2024). A primary function of boundary elements/insulators in *Drosophila* is the subdivision of the chromosome into a series of looped domains, or TADs. TAD formation is thought to depend upon the physical pairing of neighboring boundary elements. In most cases that have been examined in detail, these pairing interactions are orientation-dependent (Fujioka et al. 2016; Kyrchanova et al. 2008). Orientation dependence determines the topology of the TAD. When boundaries pair with their neighbors head-to-head, a circle-loop is generated, while head-to-tail pairing generates a stem-loop (Fujioka et al. 2016). Since *homie* and *nhomie* pair with each other head-to-tail, the *eve* TAD is a stem-loop. As we have shown (Ke et al. 2024), this loop topology enhances the physical isolation of the *eve* TAD from its neighbors. A second function of fly boundaries is mediating the pairing of homologous chromosomes, and this depends upon their ability to self-pair (Viets et al. 2019). While pairing between *homie* and *nhomie* is head-to-tail, *homie* and *nhomie* self-pairing is head-to-head (Fujioka et al. 2016), like that of other fly boundaries that have been tested (Kyrchanova et al. 2008). This orientation preference is thought to be important for mediating the juxtapositioning and precise alignment of homologous chromosomes, and it has been shown to facilitate transvection (regulatory cross-talk) between paired homologs (Fujioka et al. 2016; Lim et al. 2018), a phenomenon that is widespread in *Drosophila*.

As has been observed for the *gypsy* insulator and for boundaries from the BX-C (Geyer et al. 1990; Kravchenko et al. 2005; Li et al. 2011; Muller et al. 1999; Sigrist and Pirrotta 1997; Vazquez et al. 2006), both *homie* and *nhomie* can also mediate long-distance (100 kb to Mb) regulatory interactions (Bing et al. 2024; Chen et al. 2018; Fujioka et al. 2016; Fujioka et al. 2009). As an example, when a *homie*-containing reporter transgene is inserted at an attP site located in the *hebe* gene, 142 kb upstream of the *eve* gene, enhancers in the *eve* TAD can drive reporter expression. Reporter activation depends on the orientation of the *homie* element relative to the reporter in the transgene (e.g., eZ-CDEF in Figure 1c vs. eZ-FEDC, not shown), but does not depend on the orientation of the transgene in the chromosome (Z5 vs. 3Z in Figure 1c, d). This constraint arises because *homie* in the transgene pairs with *homie* in the *eve* locus head-to-head and with *eve*-locus *nhomie* head-to-tail (Bing et al. 2024; Fujioka et al. 2016). This orientation-specific pairing can put a reporter gene in either a favorable or an unfavorable position for the *eve* enhancers to access a transgenic promoter (Fujioka et al. 2016).

Here, we have undertaken a functional dissection of *homie*, focusing on three centrally important activities of this class of elements: long-range pairing (LR pairing), enhancer blocking, and PRE blocking. Detailed slicing and dicing of *homie* reveals a general correlation between the three activities. However, the correlation is not perfect, indicating that while the activities are related, there are some mechanistic differences. An extensive comparison of sub-element combinations provides clear cases of divergence between the requirements for each of the three insulator functions. Our results have implications for the mechanistic connections between these seemingly disparate activities.

## Results

### Multiple insulator sub-elements contribute to long-range pairing

In previous studies on long-range pairing interactions, we used a “minimal” *homie*, CDEF, which consisted of four contiguous ∼100 bp sub-elements, as it appeared to have nearly full activity (Bing et al. 2024; Chen et al. 2018; Fujioka et al. 2016). To better understand how these *homie* sequences contribute to long-range pairing, we examined the long-distance pairing activity of different combinations of sub-elements from a larger 800 bp fragment, ABCDEFGH (Figure 1b), that was previously identified as the “*homie”* insulator (Fujioka et al. 2009). According to Flybase (Gramates et al. 2022), this larger fragment extends from just downstream of the *eve* PRE to just beyond the first exon of the *TER94-RD* transcript, and it spans a DNase I hypersensitive site (Sackerson et al. 1999), as seen in boundaries from the BX-C. To test for long-range (LR) pairing interactions, we inserted *homie*-containing reporter gene constructs into an attP site in the 1^st^ exon of the *hebe* gene, 142 kb upstream of *eve* (Fujioka et al. 2016; Fujioka et al. 2009). For this analysis, we used either the single reporter transgene shown in Figure 1c or the dual reporter transgene shown in Figure 1d.

When *homie* or its derivatives have LR pairing activity, and when the insulator is in the “correct” orientation relative to the *lacZ* reporter in the transgene (e.g., eZ-CDEF in Figure 1c, but not eZ-FEDC, not shown), *lacZ* is subject to regulation by enhancers in the *eve* TAD and is expressed in an *eve-*like pattern (e.g., CDEF in Figures 2a and S1a, b). For example, *homie* CDEF in the reporter genes diagrammed in Figure 1c and d pairs with CDEF in the *eve* locus (diagrammed in Figure 1b) head-to-head (CDEF-CDEF). When this happens, the *lacZ* reporter is placed in proximity to the *eve* enhancers, while the *GFP* reporter is placed away from them, on the opposite side (in 3D space) of the paired insulators, transgene CDEF and endogenous CDEF. This results in *lacZ* being expressed in the *eve* pattern, while *GFP* is expressed very little (Fujioka et al. 2016).

**Figure 2.**
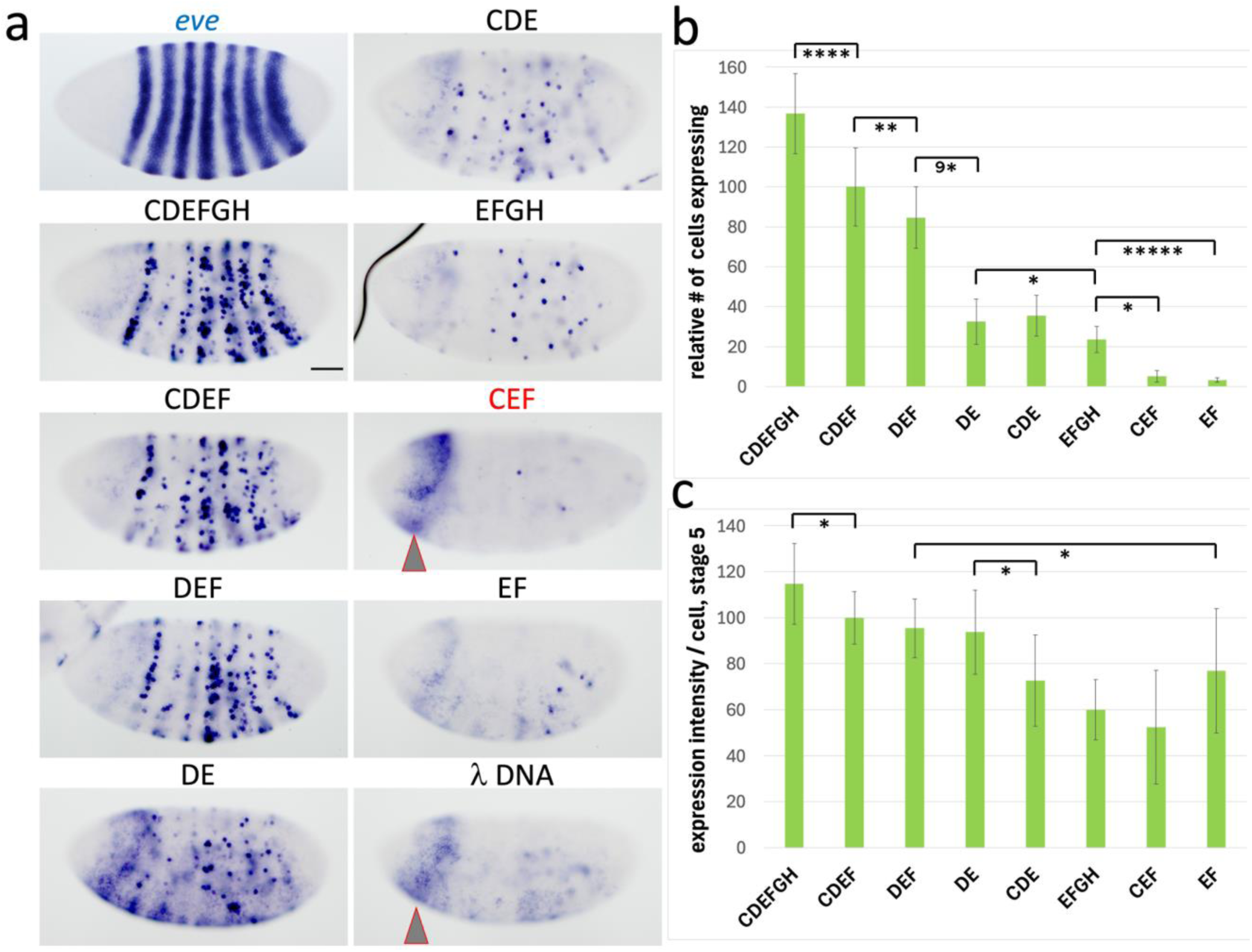
Functional dissection of the LR pairing activity of *homie*. The previously described 800 bp “full- length” *homie* (Fujioka et al. 2009) was further dissected, by dividing it into roughly 100 bp segments (A–H, see Figure 1b) and testing them in our long-range (LR) interaction assay. **a: expression from transgenic reporters at embryonic stage 5.** For reference, expression of endogenous *eve* RNA is shown at the left in (“*eve*”). For each tested region (see map in Figure 1b), *lacZ* expression from eZ-vector transgenes carrying the indicated *homie* derivative or negative control (500 bp of phage λ DNA) inserted at –142 kb is shown. The color of the label above each image indicates the transgene orientation in the chromosome: black is Z5, red is 3Z (see maps in Figure 1c). λ DNA: . Gray arrowhead indicates “head stripe” of background expression (see text). Scale bar (in CDEFGH) = 50μm. **b, c: quantification of LR pairing activity.** Images like those in **a** were analyzed (as described in Materials and Methods) to give both the number of cells expressing the reporter (*lacZ*) in an *eve* pattern (b) and the intensity of that expression in each cell (c). These quantities (number of cells expressing per embryo and average intensity of that expression per embryo) were then standardized by scaling them (linearly) relative to those of CDEF (set at 100) and λ DNA (set at 0). Graphs of averages of these quantities are shown, +/– standard deviations as error bars. The results of pairwise statistical comparisons (t-tests as implemented by Microsoft Excel) are shown as brackets connecting key pairs (see Materials and Methods for more details). Significance of the difference is indicated by the number of asterisks within each bracket: * = p < .05, ** = p < .01, **** = p < .0001, ***** = p < .00001, 9* = p < 10^−9^.

By contrast, no *eve*-like expression is seen when the same reporter transgene has an equal-length DNA segment from phage *Lambda* (“λ DNA” in Figures 2a and S1a, b). However, there is some non-*eve* related expression. Near-ubiquitous expression occurs early, but fades away by early stage 5 (“λ DNA” in Figure 2a). At stage 4, embryos also show spotty expression in yolk cells, which lasts to early stage 5 (Figure S2, olive arrowhead with black outline). At stage 5, a head stripe located anterior to *eve* stripe 1 is seen, which is more prominent with the single reporter construct (Figures 2a and S1a, b: “λ DNA”, gray arrowhead with red outline). At stage 9/10, there is some expression laterally, ventral to *eve* mesodermal expression (Figure S2, sky blue arrowhead with black outline), which is more prominent in the 3Z orientation. Some of these non- *eve*-related patterns are also seen, with varying intensities, in some of the *homie* derivatives we analyzed (e.g., the head stripe is seen with varying intensity in a number of constructs; Figure S1: gray arrowhead with red outline). We note that MicroC analysis showed a weak interaction between the λ DNA transgene and the *eve* locus, although this did not result in any detectable *eve*-like *lacZ* expression (Bing et al. 2024). Also, as described further below in the section on enhancer blocking activity, a *hebe* midline enhancer is located upstream of the attP site at –142 kb (see map in Figure 1d for the location). When *homie* is interposed between this enhancer and the transgene reporter, it blocks the enhancer from activating the reporter (for example, CDEFGH and CDEF in Figure S1a, which at stage 13 show only *eve*-like CNS expression on the ventral side). However, with λ DNA in place of *homie*, the *hebe* midline enhancer activates both the *lacZ* and *GFP* reporters (black arrowheads in λ DNA, Figures S1a and S2).

We compared the LR pairing activity of different constructs using two criteria: the number of the cells expressing the *lacZ* reporter gene in an *eve* pattern (Figure 2b) and the level of expression in these cells (Figure 2c), as determined by quantifying the average intensity of staining per expressing cell in individual embryos (see Materials and Methods for more detail). These two measures generally correlated positively with each other (with some exceptions, see below). We first extended the minimal CDEF *homie* endpoint by ∼200 bp downstream, into the beginning of the *TER94-RD* transcription unit, to give CDEFGH. The addition of the GH sub-region increased the LR pairing frequency and/or stability. This is reflected in a greater number of cells expressing the reporter at early stages (compare CDEF with CDEFGH in Figure 2a, b; see also Figure S1a), and in stronger expression at later stages, most noticeably in the mesoderm (Figure S3, left column) and CNS (Figure S3, right column) of stage 13 embryos.

Next, we assayed the LR pairing activity of the four possible combinations of three different CDEF sub-elements. Of these sub-element combinations, only DEF has LR pairing activity comparable to that of CDEF. There is a significant difference between CDEF and DEF with regard to the number of cells expressing, but not the intensity of expression (Figure 2b, c). There may be weaker expression with DEF in the mesoderm and the CNS, but this difference is small and somewhat variable (Figure S1a, b, stages 11/12 and 13, and Figure S3). Sub-element F contributes to LR pairing more than does C, since expression with DEF is stronger, especially at later stages, and seen in more cells, than is expression with CDE (Figures 2a, b and S1b). CEF shows very little stripe expression and has only weak anal plate ring (APR) expression (Figures 2a, b and S1b, red arrowhead in CEF), while CDF expression is similar to that with λ DNA, showing no *eve*-like expression (Figure S1b). Therefore, both D and E are required for LR pairing activity to generate *eve*-like expression in stripes, while enough activity remains in the absence of D, but not E, for some APR expression. We note that APR expression has been found in previous studies to be the most persistent aspect of *eve* expression produced by LR pairing, when either pairing activity is reduced or the intervening distance is increased (Fujioka et al. 2016; Fujioka et al. 2009).

Since DEF has significantly more LR pairing activity than the other tripartite combinations, we tested the relative contributions of its three sub-elements, DE, EF, and DF. As shown in Figures 2 and S1c, DF showed no clear evidence of LR pairing activity. While EF rarely showed stripe expression at stages 5-8, it did give APR expression at stage 11/12 (S1c: red arrowhead in EF). DE, on the other hand, clearly has more LR pairing activity than either DF or EF. In every DE embryo, a small number of cells expressed *lacZ* at stages 5-8 (Figures 2 and S1c), as well as in the APR (stages 11-13, Figure S1c: red arrowheads in DE) and CNS (stage 11/12, Figure S1c: yellow arrowhead in DE). However, as is evident from the large number of cells expressing *lacZ* in stage 5-8 DEF embryos (Figure 2b), the F sub-element clearly bolsters the LR pairing activity of the DE combination (Figures 2a, b and S1b, c, compare DEF with DE). None of the single sub-regions (D, E, or F alone) gave any *eve*-like expression (Figure S1d). We note that DE, DF, D, E, and F gave broad, “background” expression at stage 5 similar to that of the λ DNA control (Figures 2a and S1). Interestingly, this phenomenon may be analogous to the background expression that is often more prominent when small enhancer fragments that retain little or no “specific” activity are tested in reporter transgenes. Perhaps strong insulators, like strong enhancers, tend to harbor some repressive activity. Alternatively, the absence of background expression with our stronger insulator fragments may be a consequence of their stronger enhancer blocking activity.

To further test the contribution of the GH region to LR pairing activity, we combined GH with EF (EFGH). As shown above, EF on its own has almost no LR pairing activity, except weak APR expression at stage 11/12 (Figure S1c, red arrowhead in EF). In contrast, a spotty *eve*-like pattern of *lacZ* expression is observed with EFGH throughout embryogenesis: *lacZ* is expressed in more cells than with EF at stages 5-8 (Figures 2a, b and S1a), in the APR at stages 11-13 (Figure S1a, red arrowheads), and in the mesoderm at stage 13 (Figure S1a, green arrowhead). However, the intensity of expression is weaker than with either CDEF or DEF, and far fewer cells express the reporter gene (Figures 2 and S1a, b), indicating that GH can only partially substitute for the D sub-element.

On the other hand, the AB sub-element combination cannot replace EF, as ABCD has no LR pairing activity (Figure S1a). In other experiments, we further tested the AB region by creating ABDE in the context of the double reporter transgene (Figures 3 and S1e). The AB region does not increase the LR pairing activity of DE, since neither the average number of expressing cells nor the average intensity of reporter expression in an *eve* pattern with ABDE is significantly increased over that with DE alone.

**Figure 3.**
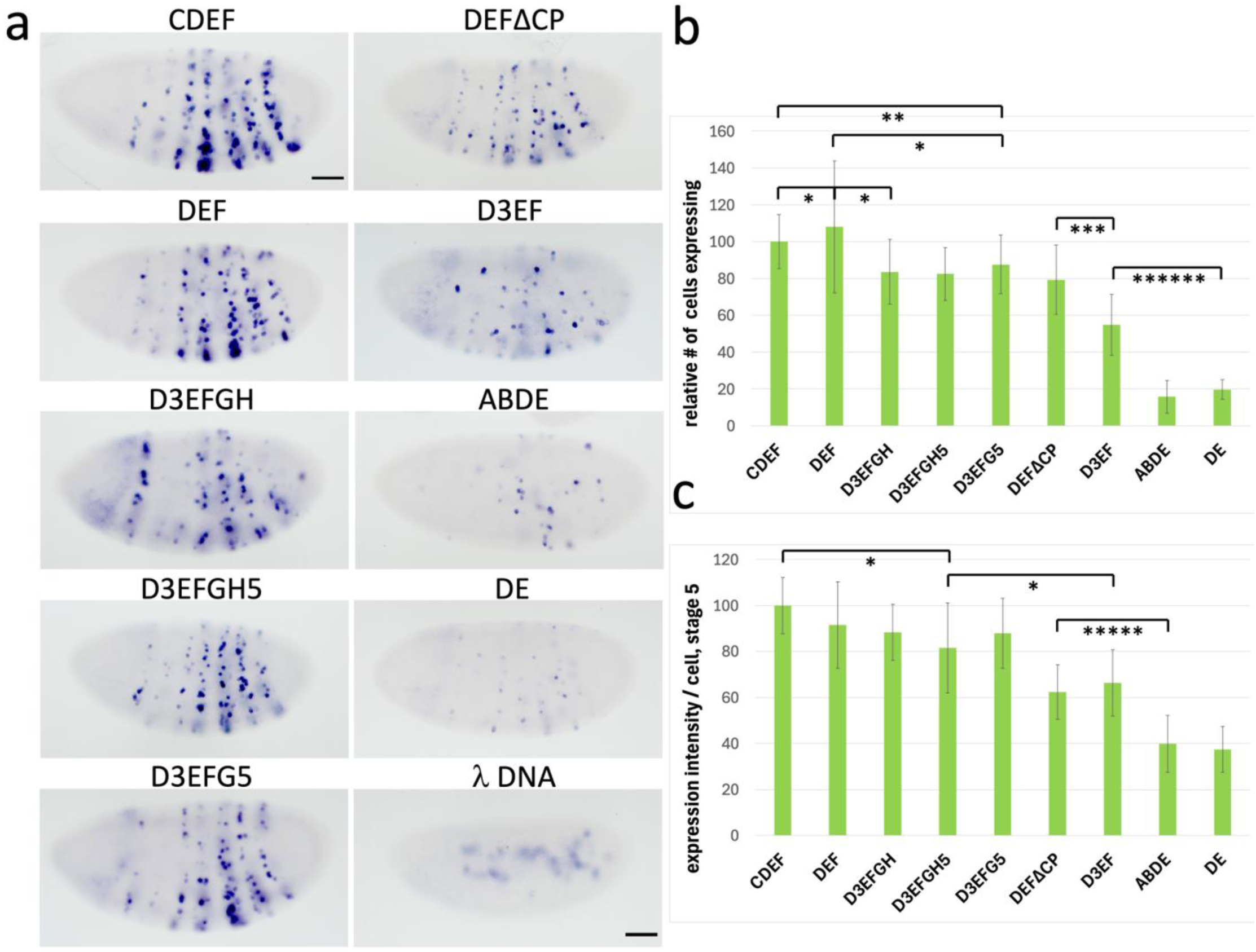
Quantitative comparison of the LR pairing activities of *homie* derivatives. **a:** transgenic reporter (*lacZ*) expression is shown from the indicated eZ-eG transgenes in the Z5 orientation at embryonic stage 5. In D3EF, the 5’ (left) half of D was removed from DEF (the Su(Hw) site is still present), while in D3FE, the orientation of section EF was additionally reversed. In D3EFG5, D3EF is extended to just downstream of the *TER94-RA* start site; in D3EFGH5, it is extended to just downstream of the *CR45324* start site; in D3EFGH, it is extended through the end of *homie* subregion H, which includes the *TER94-RD* start site. Scale bar (in “CDEF”) = 50μm. **b, c:** Images like those in **a** were quantified and graphed as described in Figure 2b, c. Significance of key differences is indicated by the number of asterisks within each bracket: * = p < .05, ** = p < .01, *** = p < .001, ***** = p < .00001, ****** = p < 10^−6^.

### Insulator protein binding sites are required for long-range pairing

The experiments in the previous section show that DE harbors more of *homie*’s LR pairing activity than any other 2-sub-element combination. Genome-wide analysis of the distributions of the Su(Hw) and Cp190 proteins showed that both proteins localize to *homie* and *nhomie* (Baxley et al. 2017; Cuartero et al. 2014; Negre et al. 2010; Schwartz et al. 2012; Soshnev et al. 2012; Wood et al. 2011). Sequence-searching of *homie* identified a Su(Hw) binding site motif (Baxley et al. 2017; Negre et al. 2010) in D, and a Cp190- associated sequence motif (Cuartero et al. 2014) in E. We tested whether these sites are involved in LR pairing. We mutated the Su(Hw) site in the context of DEF (DEF-ΔSu(Hw), or DEFΔSH for short). As shown in Figures 4 and S5, mutating the Su(Hw) site results in a loss of most of the LR pairing activity; however, weak APR expression remains (Figure S5, red arrowheads). Interestingly, the *gypsy* insulator, which has multiple Su(Hw) binding sites, did not show any LR pairing activity (Figure S1e), indicating that Su(Hw) binding sites are not sufficient for this activity. Mutating the Cp190-associated sequence in the context of DEF reduced LR pairing (DEF-ΔCp190, or DEFΔCP for short, Figures 3 and S5), although the effect is not as drastic as that of the Su(Hw) site mutation (Figure S5). We note that this Cp190-associated sequence motif from Cuartero et.al. is similar to a Pita binding motif reported by Maksimenko, *et al*. (Maksimenko et al. 2015). That study reported that the Pita motif is similar to the Cp190 motif identified by Schwartz, *et al*. (Schwartz et al. 2012), and that Pita is capable of recruiting Cp190 (see Figure S4 for excerpts of genome-wide ChIP data).

**Figure 4.**
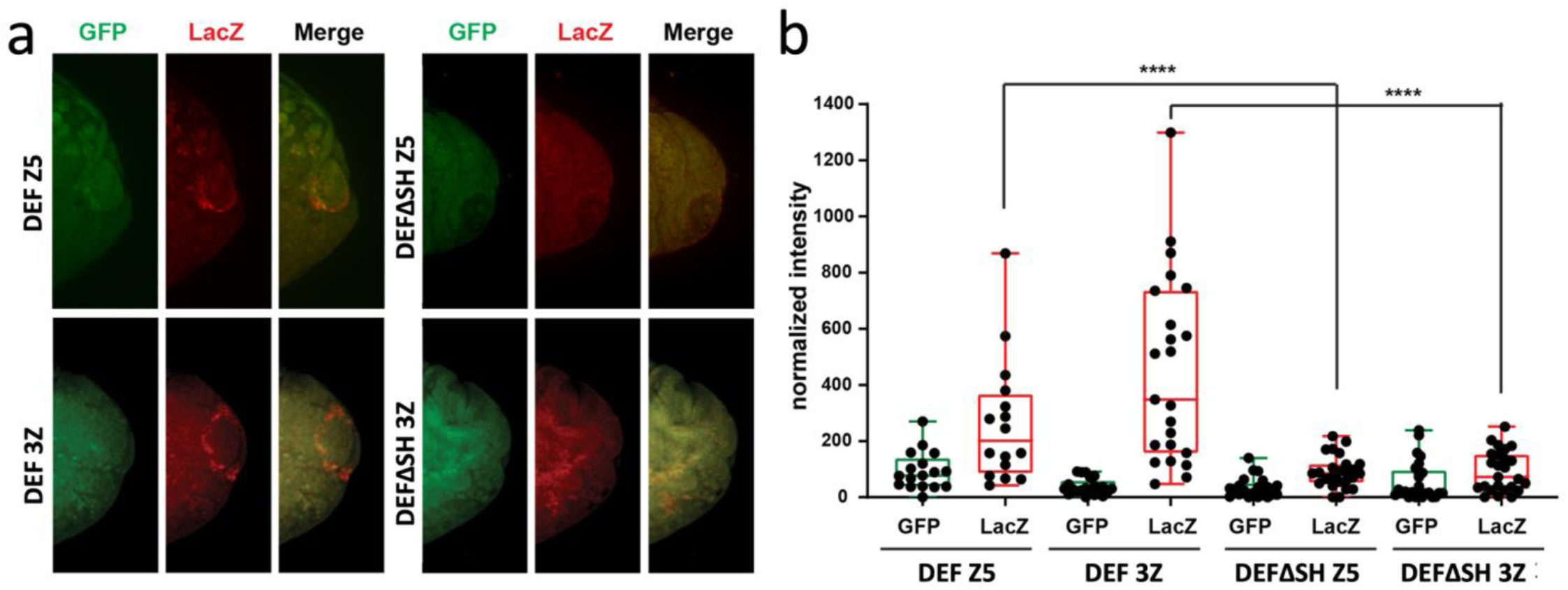
The one consensus binding site for Su(Hw) in *homie* contributes strongly to long-range pairing. smFISH analysis was performed on stage 15 DEF and DEFΔSH eZ-eG transgene embryos. Expression of *lacZ* and *GFP* in the anal plate region is shown in **a**. Quantification of *lacZ* expression is shown in **b** (**** indicates significance of the difference at the p < 0.0001 level). Each image is 80μm in width.

To further quantify the effects of the Su(Hw) site mutation, we used smFISH on stage 15 embryos. In the experiments of Figure 4, DEF and DEFΔSH dual-reporter transgenes were inserted into the attP site at –142 kb in each orientation, Z5 and 3Z (see diagram in Figure 1d). We then probed for *lacZ* and *GFP* expression driven by the APR enhancer. We focused on this LR activity because expression is less stochastic in this tissue than in other expressing tissues, and most *eve*-expressing APR cells also show reporter expression in the starting DEF construct. In both the Z5 and 3Z transgene orientations, *eve*-like *lacZ* expression is observed in the APR, while *GFP* expression is only rarely detected (Figure 4). This is consistent with the digoxigenin staining (Figure S5 for *lacZ* and Figure S7 for *GFP*). Quantitation of *lacZ* mRNA in the APR showed that mutation of the Su(Hw) recognition sequence in the D sub-element very significantly impairs LR pairing activity, resulting in a substantial drop in *lacZ* expression (Figure 4b). This is true when the transgene is inserted in either orientation, 3Z or Z5.

Next, we tested whether the Su(Hw) site is fully responsible for the activity of D, using our dual reporter transgene. Here, we added back the 3’ half of D (“D3”), which contains the Su(Hw) site, to EF, which has only very weak LR pairing activity (see above), to make D3EF. The addition of the D3 fragment clearly restores some LR pairing activity, since the *lacZ* reporter is expressed in an *eve* pattern throughout embryogenesis (Figures 3 and S5). However, the activity of D3EF is significantly weaker than that of DEF (Figures 3 and S5), suggesting that sequences in the 5’ (left) half of D interact with a factor(s) that contributes to LR pairing. On the other hand, the activity of D3EF is significantly stronger than that of DE (Figure 3), indicating that the F region can more than substitute for the 5’ portion of D.

Finally, we tested whether the order of sequences within *homie* is important for its activity. To do this, we reversed the order of EF in D3EF, to make D3FE. As shown in Figure S4, changing the order abolishes LR pairing activity, as the *eve* enhancers fail to activate *lacZ* expression. It seemed possible that the change in the order of the sub-elements altered the orientation dependence of D3EF. That is, if the orientation dependence were due to EF alone, then D3FE would align with FE of the endogenous *homie* (see Figure 1b). In this case, the *lacZ* reporter would be placed away from the *eve* enhancers (and not be expressed). Since D3FE is inserted in the dual reporter, we were able to test this possibility by assaying *GFP* expression. Figure S7 shows that GFP is also not expressed in the D3FE dual reporter, indicating that LR pairing activity has indeed been lost with the change in sub-element order, not switched in its orientation preference.

Consistent with the maintenance of orientation specificity with these constructs whenever there is significant LR pairing, we found that neither of the other dual reporter constructs tested (DEF-ΔSu(Hw) and DEF) detectably expressed *GFP* in an *eve* pattern (Figure S7; however, they do express *GFP* in the pattern of the *hebe* midline enhancer at stage 13, since the enhancer is not able to be shielded from the *GFP* reporter in this orientation of the transgene).

### Contribution of *TER94*-associated elements to LR pairing activity

In the experiments described above, GH contributed to LR pairing activity in the context of both CDEF and EF *homie*. G contains the 5’-most *TER94* transcription start site (which produces transcript *TER94-RA*; *TER94* is transcribed away from the *eve* locus), while H contains a non-coding lncRNA start site (CR45324, which is transcribed toward the *eve* locus) and the initiation site for transcript *TER94-RD* (Flybase, Gramates et al. 2022). Because D3EF showed weak but clear-cut LR pairing activity (Figures 3, S5, and S6), we tested whether the adjacent GH region enhances the LR pairing activity of D3EF. Indeed, D3EFGH showed considerably stronger activity than does D3EF (Figures 3 and S6). We then trimmed back the added region to exclude the start sites for both the non-coding RNA and *TER94-RD*. This resulted in no apparent loss of LR pairing activity (D3EFGH5, Figures 3 and S6). We further trimmed it to just downstream of the *TER94-RA* start site, again with no apparent loss of activity (D3EFG5, Figures 3 and S6). This suggests that it is the ∼50 bp region just downstream of the F region that harbors most, if not all, of the LR pairing activity of GH. Since this facilitating region includes the *TER94-RA* start site (to +2 nt), it may contain a basal promoter element from this housekeeping gene. This increased activity is still a bit less than that of CDEF (Figures 3 and S6). It is also worth noting our previous finding that *TER94* promoter activity is enhanced by sequences between the first and third exons of the *TER94-RA* transcript (Fujioka et al. 2013), which are mostly not included in this insulator-facilitating fragment. So, the sequences facilitating *homie*’s insulating activity and those responsible for *TER94*’s transcriptional activity seem to be mostly, if not entirely, separable, with the caveat that we can’t rule out a minor contribution of *homie*’s 3’ end (the 5’ portion of region G) to the level of *TER94* expression.

### Enhancer blocking activity only roughly correlates with other insulator activities

Next, we investigated how LR pairing activity correlates with enhancer blocking activity. As an assay, we tested the ability of different *homie* sub-elements to block the *hebe* enhancer from activating the *lacZ* transgene reporter (Fujioka et al. 2016). This enhancer is located upstream of the –142 kb attP site in a part of the first intron of the *hebe* gene (Figure S8a, *hebe*-2). Interestingly, the *hebe* intron also contains a weak APR enhancer. However, it is active only after stage 16 (Figure S8a, *hebe*-1), unlike the *eve* APR enhancer.

Because of this, we assessed *eve*-like APR expression only at stages 11-15. Consistent with these enhancer activities, both expression patterns were reported by the Berkeley Drosophila Genome Project (Tomancak et al. 2007). The 3^rd^ activity we identified, driving expression in the gut (Figure S8a, *hebe*-3), was not reported by the Genome Project, and we did not see this expression from our λ DNA transgene (Figures S1a, b and S2). It is possible that the gut expression driven by *hebe*-3 depends on something in the chromosomal environment of the attP site we used for this enhancer mapping. This activity is not relevant to our assay.

The relative position of the midline enhancer to the transgene is shown in Figure 1d. In the Z5 orientation, when CDEF is located between the *lacZ* reporter and the *hebe* enhancer, it blocks the enhancer from activating reporter expression, and only *eve*-like expression is seen in the CNS (Figure S8b, CDEF and all the other insulators in the left column except CDE; see Figure S3 for details of *eve*-like expression). On the other hand, λ DNA has little or no blocking activity, and so the *hebe* enhancer is able to activate *lacZ* expression in CNS midline cell clusters, mimicking *hebe* expression at this stage (“λ DNA”, Figures S1a, b and S8b: in S8b, this is also true, to varying degrees, for those in the middle and right columns, and for CDE in the left column). In the 3Z orientation, the midline enhancer can activate *lacZ* expression independent of the presence of *homie*, since *homie* is not located between the enhancer and the *lacZ* reporter (see Figure 1d for a diagram, and CDEF, CEF, and λ DNA in Figure S1c).

Enhancer blocking activity was tested at the two stages of embryogenesis when the *hebe* enhancer is active in cells in the CNS midline (Figure S8a, b, *hebe*-2, late stage 12 and stage 13, persisting into stage 16). One set of fragments was tested in the context of the single reporter transgene (the eZ vector, Figures 5a and S8b), and another, overlapping set was tested in the double reporter transgene (the eZ-eG vector, Figures 5b and S8b). The two assays gave a consistent order of activity with all of the constructs common to the two assays (CDEF, DEF, DE, and λ DNA). However, the apparent strength of the enhancer blocking activity of DE in the eZ-eG vector is less than that seen with the eZ vector. This difference in apparent blocking strength allowed us to make clear distinctions using the eZ-eG vector between the activities of the five constructs whose activities lie between those of DEF and DE (Figure 5b).

**Figure 5.**
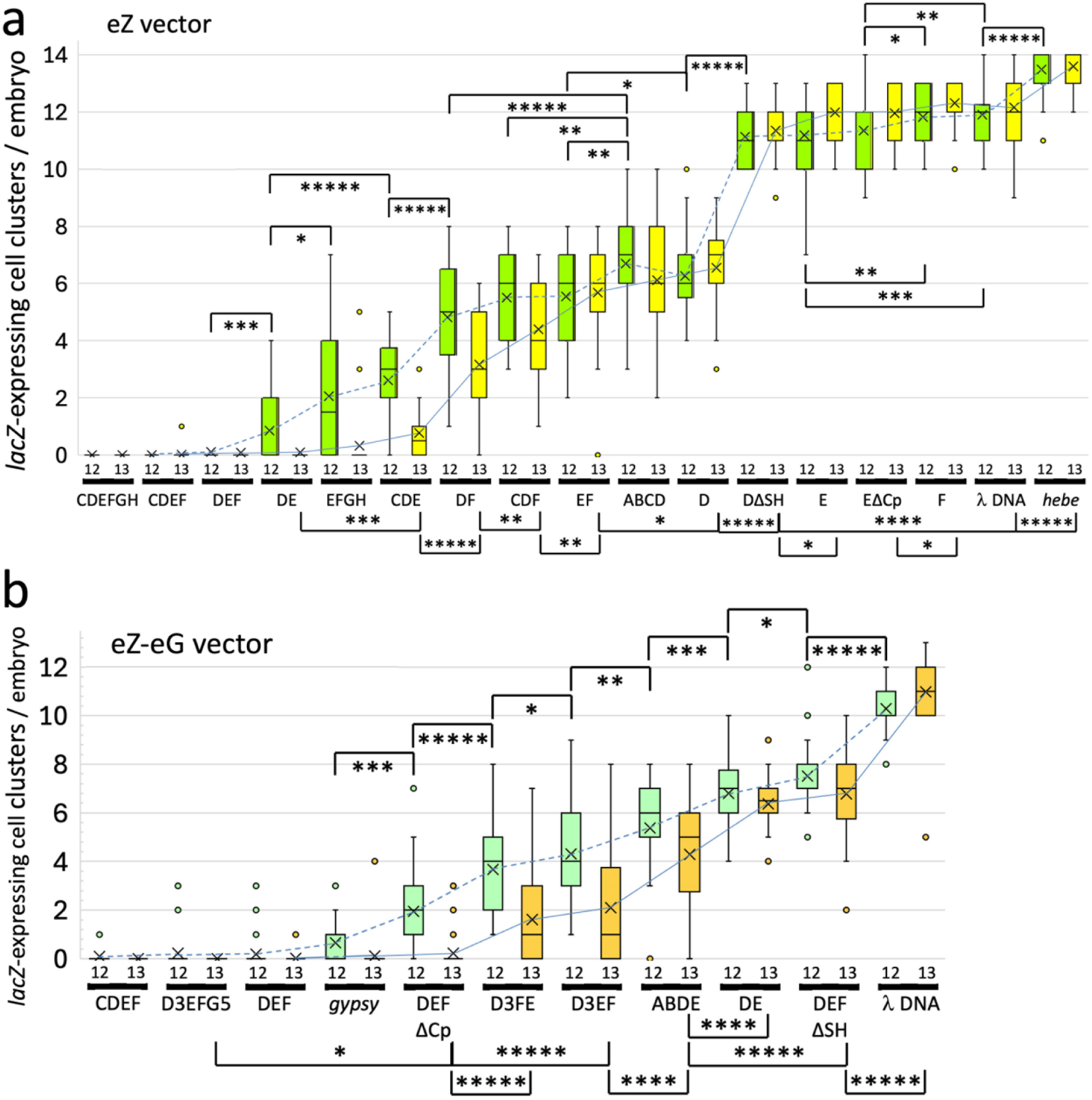
Enhancer blocking activity of *homie* and its derivatives. Box-and-whiskers plots of the average and distribution of the number of CNS midline cell clusters expressing the *lacZ* reporter at late embryonic stages 12 and 13. Each box represents the 25^th^ to 75^th^ percentile range. The horizontal line and “X” mark the median and average, respectively. Whiskers represent non-outlier data points, and open circles represent outliers. The averages for each transgene at stage 12 and at stage 13 are connected with dotted and solid lines, respectively. Representative CNS images and the number of embryos counted for each transgene are shown in Figure S5b. Those with a higher number of visible clusters (which is plotted along the y-axis) have less enhancer blocking activity. The *homie* sub-elements present in the transgene are listed below each pair of plots: names are as in the text; e.g., “ΔSH” has the Su(Hw) consensus binding site mutated, and “ΔCp” has the Cp190-associated site mutated. Constructs were ordered from left to right based on the average number of midline cell clusters expressing. The results of pairwise statistical comparisons (t-tests as implemented by Microsoft Excel) are shown as brackets connecting each pair. Stage 12 comparisons are shown above the plots, while those at stage 13 are shown below the plots. Significance of the difference is indicated by the number of asterisks within each bracket: * = p < .05, ** = p < .01, *** = p < .001, **** = p < .0001, *****= p < .00001. **a:** eZ vector transgenes. **b:** eZ-eG vector transgenes.

For the tripartite combinations DEF, CDE, and CDF, blocking activity follows a pattern roughly similar to that of LR pairing activity (Figures 2, S1b, 5a, and S8b). DEF is the most effective enhancer blocker, followed by CDE, and then CDF. Because we obtained a transgenic line carrying only one of the two possible orientations of CEF (the 3Z orientation, in which the insulator fragment is not between the reporter and the *hebe* midline enhancer), we were not able to determine its enhancer blocking activity. For the three 2- element combinations from DEF (DE, DF, and EF) in the eZ vector, DE is almost as effective in stage 13 embryos as DEF, while DF and EF each have considerably weaker activity (Figures 5a and S8b), roughly paralleling their LR pairing activities. The addition of GH to EF (EFGH) increased enhancer blocking activity (Figure 5a), similar to the effect seen on LR pairing (Figures 2a, b and S1a, c). Although we did not see an increase in LR pairing activity of ABDE over DE (Figures 3 and S1e), addition of AB significantly increased the enhancer blocking activity of DE (Figure 5b).

Since the Su(Hw) and Cp190 sites contribute to the LR pairing activity of DEF (Figures 3, 4, and S5), we tested the effects of mutations in these recognition sequences (Figures 5b and S8b). As was the case for LR pairing activity, mutation of the Cp190 site had a modest effect on DEF’s enhancer blocking activity (at stage 12, but not at stage 13). In contrast, mutating the Su(Hw) site led to a substantial loss in this blocking activity. Finally, we assayed the enhancer blocking activity of individual sub-elements (D, E, and F), as well as the effects of mutating the Su(Hw) and Cp190 sites, located in fragments D and E, respectively (Figure 5a). D alone showed substantial activity, while E alone gave a low but detectable activity, and F alone showed no significant difference from λ DNA. When the Su(Hw) site was mutated, it abolished activity in the context of D alone. Mutating the Cp190-associated site has no clear effect on the activity of E, as a low activity is retained in EΔCP (Figure 5a). The addition of region ABC to D did not increase enhancer blocking activity (Figure 5a, ABCD), indicating that there is no significant enhancer blocking activity within ABC. Again, these effects on enhancer blocking activity parallel the effects of the same alterations on LR pairing activity.

There are exceptions to this correlation, however. Most dramatic is the case of the *gypsy* transposon, which showed strong enhancer blocking activity (Figure 5b), but no LR pairing activity (Figure S1e). Seven of the other constructs tested in the eZ-eG vector showed less enhancer blocking activity than *gypsy*, but more LR pairing activity (see Figure 7 for a summary). There are also examples among the *homie* derivatives. EF has detectable LR pairing activity (Figures 2 and S1c), whereas DF does not (Figure S1c), while DF has significantly more enhancer blocking activity at stage 13 than does EF (p < 4.7 x 10^−8^, Figure 5a). Another example is provided by a comparison of the two *homie* derivatives D3EF and D3FE (in which the order of the EF region is reversed; see Figure S5a for a diagram). While D3FE is unable to mediate LR pairing with the *eve* TAD, D3EF did show LR pairing activity (Figures 3 and S5). In contrast, both showed modest enhancer blocking capability (Figure 5b; in fact, at stage 12, D3FE showed significantly stronger blocking activity than D3EF, p < 0.03). This suggests that, at least from the location of the *hebe* locus, LR pairing activity is much more sensitive to the order of binding sites for insulator proteins than is enhancer blocking activity. In summary, the correlation is not perfect for all constructs, suggesting that while the two activities are closely related, there are some mechanistic differences (see Discussion).

### PRE blocking activity

We showed previously that *homie* has the ability to block the spread of repressive chromatin (barrier activity), from an *eve* pseudo-locus transgene into the neighboring gene, *TER94*. This spreading depends on the *homie*-adjacent *eve* PRE (Fujioka et al. 2013). *TER94* promoter activity is repressed when *homie* is removed, and this correlates with the spreading of the histone modification H3K27me3, characteristic of PcG-dependent repressive chromatin. This loss of expression is not due to reduced *TER94* promoter activity caused by removing *homie*, since removing the *eve* PRE in addition to *homie* fully restores expression (Fujioka et al. 2013). In this transgenic context, when *homie* is present, *TER94* promoter-driven GFP expression is high in ovaries dissected from adult females (Figure S9a, wt), while *TER94*-driven GFP is repressed when *homie* is removed (Figure S9a, Δ*homie*).

Whether full-length *homie* (ABCDEF) is replaced by CDEF or by any of the sub-element combinations DEF, CDE, CEF, CDF, DE, DF, or EF, *TER94* promoter-driven GFP expression is undiminished, indicating that PRE blocking is still complete (Figure S9a, b). As shown in Figure 6a, we also tested other *homie* sub-elements.

**Figure 6.**
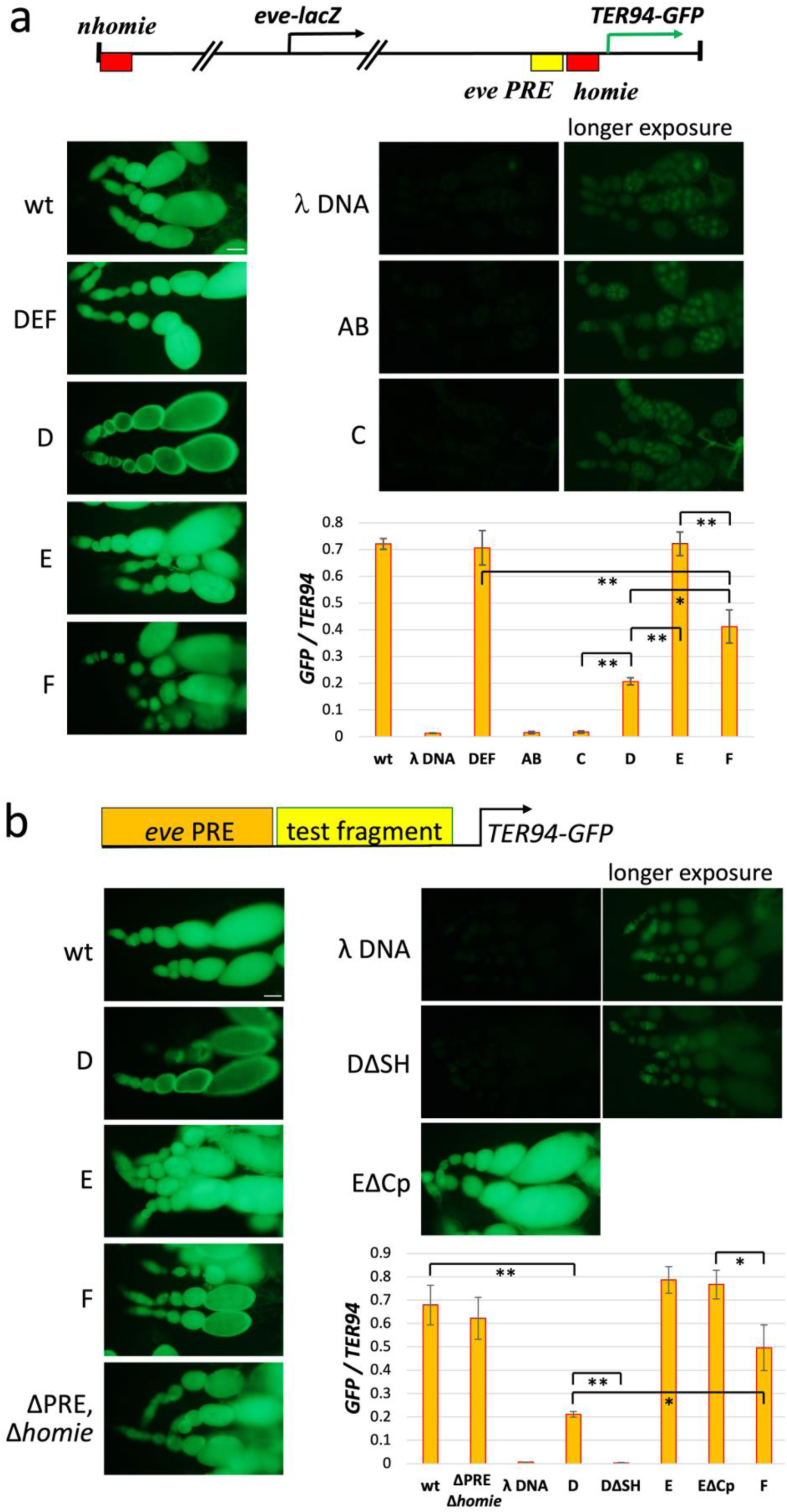
Multiple, non-overlapping *homie* sub-elements are sufficient for PRE blocking. Top: **Diagram of** the *eve* pseudo-locus transgene (Fujioka et al. 2013). **Images:** Live images of *GFP* expression in dissected ovarioles. Scale bars = 50μm. **Graphs:** quantification of *GFP* expression by RT-PCR. *GFP* RNA levels were normalized to endogenous *TER94* RNA in each sample. The averages and standard deviations of 3 independent data sets for each sample are shown. The results of pairwise statistical comparisons (t-tests as implemented by Microsoft Excel) are shown as brackets connecting each pair. Significance of the difference is indicated by the number of asterisks within each bracket: * = p < .05, ** = p < .01. **a. wt**: intact pseudo-locus inserted at 74A2 (Fujioka et al. 2013). **DEF**, **D**, **E**, **F**, **AB**, **C**: the ABCDEF region of *homie* was replaced with each of these derivatives. In each case, λ DNA was used to make the spacing between the 3’ PRE and the *homie* element similar to that in the “wt” pseudo-locus. **λ DNA**: the same sequence used in Figure 2 replaced ABCDEF. For λ DNA, AB, and C, longer exposures of the same ovaries are shown at the right. **b.** A shorter assay construct, diagrammed at the top, was used, consisting of the 3’ end of the *eve* pseudo-locus, starting just upstream of the PRE, inserted at 24B1. The ABCDEF region of *homie* was replaced with each of these derivatives, as in **a**: **wt**, **D, E, F, λ DNA**; **DΔSH, EΔCp**: the same point mutations of the Su(Hw) and Cp190- associated sites in *homie* as in Figure 3 were introduced; **ΔPRE, Δ*homie***: the *eve* PRE and ABCDEF *homie* together were replaced by 1.3 kb of phage λ DNA. For λ DNA and DΔSH, longer exposures of the same ovaries are shown at the right.

Of these, AB and C have no barrier activity, while E alone is sufficient to completely block the repressive effect of the *eve* PRE. D and F each have partial barrier activity: GFP expression is diminished, but not as much as when *homie* is replaced by λ DNA (compare “λ DNA” with D and F in Figure 6a). It is also worth noting that this barrier activity does not depend on the orientation of *homie* (Figure S9a, CDEF vs. FEDC).

Above, we showed that the Su(Hw) binding site in the D region is required for most, but not all, of the activity of D in the LR pairing assay. We asked whether the Su(Hw) binding site is also required for PRE blocking activity. This set of experiments was performed in a different context than the one used for Figures 6a and S9. Instead of the full-length *eve* locus in the transgene, we used only the 3’ end of the locus, which contains all of the essential elements for this assay; namely, the *eve* 3’ PRE, *homie*, and the *TER94* promoter driving GFP (diagrammed at the top of Figure 6b). In addition to the transgene construct being different, the insertion site of the transgene is different. Despite these differences, the results are very similar to those shown in Figure 6a (compare wt, D, E, F, and λ DNA between Figure 6a and b). As expected, when the PRE is not present in the transgene, there is no repression of *TER94*-driven GFP expression, even in the absence of any insulator sequences (“ΔPRE, Δ*homie*”, Figure 6b; here, both the PRE and *homie* are replaced by equal-length stretches of λ DNA). We tested the same mutation in this assay, and found that the Su(Hw) site is required for the D region to show any barrier activity (Figure 6b, DΔSH). We also tested the requirement for the Cp190-associated site in region E: it does not have any apparent effect on the barrier activity of E (Figure 6b, E vs. EΔCp). In summary, unlike their lack of enhancer blocking ability, both E and F alone showed PRE blocking activity, suggesting that enhancer blocking and PRE blocking activities may have partially distinct mechanisms, and/or that their mechanisms may be differentially context-dependent (see Discussion).

## Discussion

In their endogenous locations, *homie* and *nhomie* are separated by only 16 kb. However, both elements can pair with themselves and with each other when separated by multiple TADs and TAD boundaries (Bing et al. 2024; Chen et al. 2018; Fujioka et al. 2016; Fujioka et al. 2009; Ke et al. 2024). In addition to this LR pairing activity, *homie* can block enhancer-promoter interactions (Fujioka et al. 2016; Fujioka et al. 2021; Fujioka et al. 2009) and act as a barrier to the spread of PcG silencing (Fujioka et al. 2013). In the studies reported here, we have undertaken a functional dissection of the *homie* boundary and identified sequences/sub-elements that are important for these three activities. Figure 7 summarizes the order of activity of each of these elements for each of the three activities.

**Figure 7.**
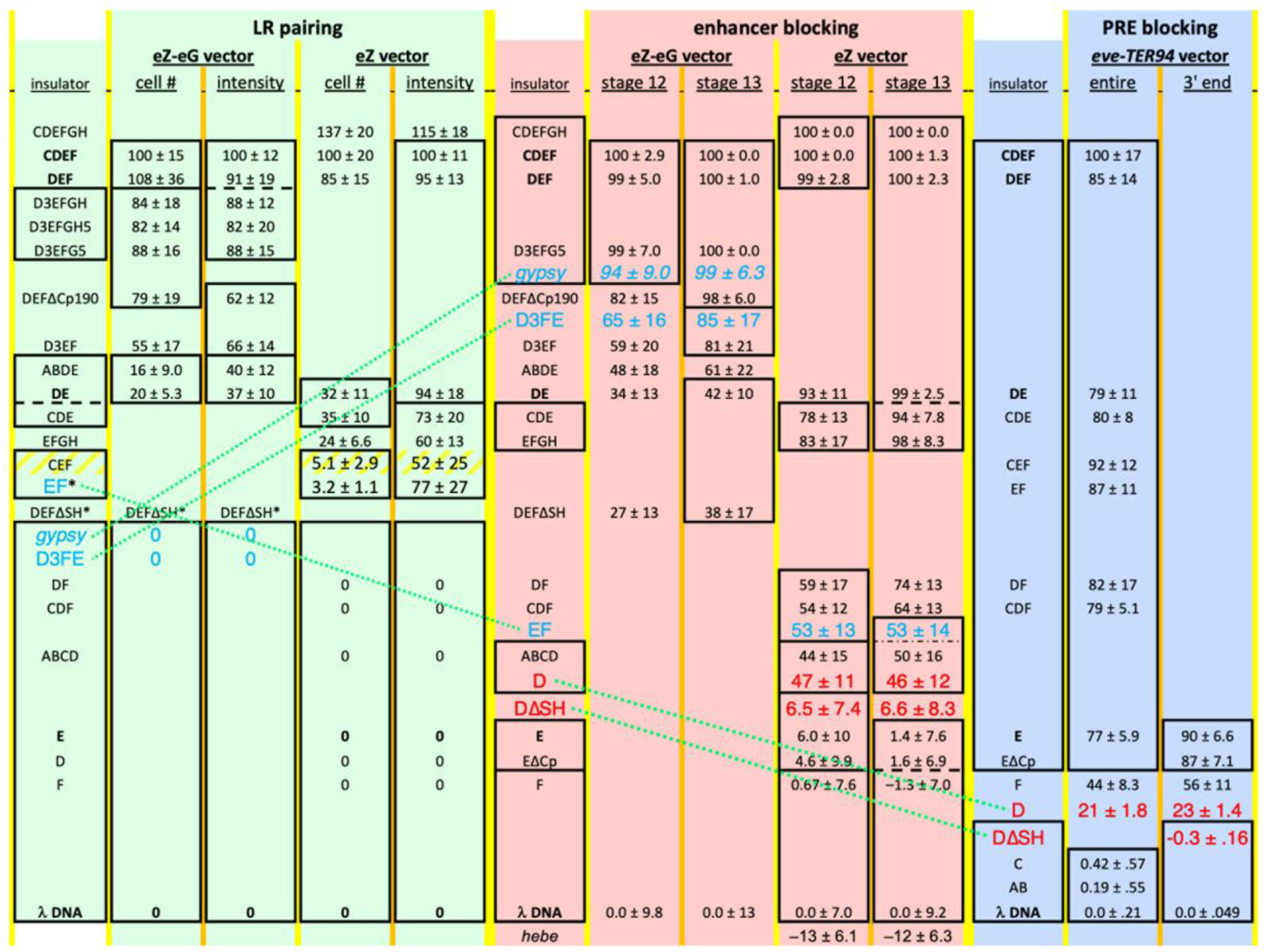
Relative strengths of LR pairing, enhancer blocking, and PRE blocking activities do not always correlate. Constructs within each column (pale green background for LR pairing, pink for enhancer blocking, and blue for PRE blocking) were ordered based on their relative levels of activity in the indicated assay. Constructs that are common to all 3 assays and both vectors are listed in boldface in the first column under each activity. Those within boxed groups in each column have activities that are not distinguishable in that assay. Averages are given of each activity (+/– standard deviations) relative to those of CDEF (set at 100) and λ DNA (set at 0), as in the graphs of Figures 2 and 3. Asterisks indicate slightly better LR pairing than those in the group just below, based on detectable APR reporter expression. Green dotted lines connect those (blue or red text) that clearly differ in their order of activity in two of the assays (see text).

### Different stringencies for different insulator activities

For LR pairing activity, DEF is our minimal element, since it gives an *eve* pattern of *lacZ* throughout embryogenesis, and the expression pattern and intensity are similar to those of CDEF. Shortening this to DE significantly weakens activity, since many fewer cells express *lacZ* (Figures 2, 3, and S1a-e). For enhancer blocking activity, in the context of the eZ vector, DE is our minimal element. Although we observe transient *hebe* enhancer-driven midline expression at stage 12, by stage 13, this expression is not seen. Removal of E to give D alone causes substantial loss of enhancer blocking activity, but considerable activity remains, while E alone shows a lower but detectable activity, and F shows no significant activity (Figures 5a and S8b). For PRE blocking, E alone has full activity, while D and F alone each have partial activity, F having more than D (Figure 6). Overall, the requirements for LR pairing activity are the most stringent, followed by enhancer blocking, with PRE blocking showing the least stringent requirements in our assays.

As noted in Results, we tested enhancer blocking activity using both the eZ and eZ-eG vectors. While the two vectors gave consistent results, in that there were no discrepancies in the relative strengths of enhancer blocking between the constructs tested in both vectors, the absolute level of enhancer blocking seen with the eZ-eG vector was lower. The two differences between them are 1) the eZ-eG vector has two reporter genes (divergently transcribed), which places *lacZ* about 1.4 kb further away from the *hebe* enhancer than it is in the eZ vector, and 2) there is a stretch of λ DNA (∼400 bp) inserted as a spacer between the smaller test fragments (such as DE) and *lacZ* in eZ that is not present in eZ-eG. While we cannot be sure of the cause of the observed quantitative difference in enhancer blocking, it is possible that the extra distance (∼1 kb) between the *hebe* enhancer and *lacZ* in the eZ-eG vector makes it easier for the enhancer to loop around the insulator sequence and activate *lacZ* in the *hebe* pattern, thus decreasing the measured enhancer blocking activity.

### LR pairing vs. enhancer blocking

We compared LR pairing activity and enhancer blocking activity, and found that the two roughly correlate (summarized in Figure 7). However, this correlation is not perfect. One clear example is provided by D3EF and D3FE, which each show substantial enhancer blocking activity (Figure 5b). However, while D3EF has clear-cut LR pairing activity (although weaker than that of DEF), D3FE has none (Figures 3, S5, and S6).

Reversing the orientation of EF relative to the D3 region likely causes a change in the order of insulator protein binding across the element. Thus, the “correct” ordering of insulator binding proteins along the chromosome may be required for LR pairing. This is readily explained by individual insulator proteins requiring specific partners for effective pairing. This also explains a preference for self-pairing, which is a property of several of the known insulator binding proteins (Ghosh et al. 2001; Vogelmann et al. 2014; Zolotarev et al. 2016). We also showed that the single direct Su(Hw) binding site is very important for LR pairing activity by *homie* (Figures 4, S5, and S7, DEF-ΔSH). In contrast, the *gypsy* insulator, which contains 12 Su(Hw) binding sites (Parkhurst et al. 1988; Spana et al. 1988) and can self-pair (Cai and Shen 2001; Muravyova et al. 2001), shows no detectable LR pairing activity with the *eve* TAD (Figure S1e). On the other hand, it has strong enhancer blocking activity in our blocking assay (Figure 5b). These observations thus suggest that additional insulator proteins besides Su(Hw) are required to form specific and stable long-range pairing interactions between copies of the *eve* insulators. Consistent with this idea, genome-wide analyses have shown that many of the known insulator binding proteins are bound at *homie* and *nhomie* (Bag et al. 2021; Baxley et al. 2017; Cuartero et al. 2014; Cubeñas-Potts et al. 2016; Li and Gilmour 2013; Li et al. 2015; Maksimenko et al. 2015; Matzat et al. 2012; Ramírez et al. 2018; Schwartz et al. 2012; Soshnev et al. 2012; Van Bortle et al. 2014; Van Bortle et al. 2012; Wood et al. 2011; Zolotarev et al. 2016).

As noted above, enhancer blocking seems less demanding than LR pairing. For example, D alone has enhancer blocking activity but no LR pairing activity. Like LR pairing, the enhancer blocking activity of D strongly depends on the Su(Hw) site (Figure 5a), indicating that Su(Hw) contributes substantially to both of these activities. Interestingly, region F alone shows no enhancer blocking activity, while E alone has a very low amount (albeit significantly above that of λ DNA), but the two together (EF) have much more activity (in fact, significantly more than that of D alone, Figure 5a). Likewise, the blocking activity of D is clearly augmented by adding E region (DE, Figure 5a). These observations suggest that, like LR pairing, a combination of proteins contributes synergistically to enhancer blocking activity. The EF region is also capable of detectable LR pairing with the *eve* locus, whereas DF and CDF are not (Figures 2 and S1b, c). This is consistent with the importance for LR pairing of the order and spacing of protein binding sites along the insulator, whereas the relative positions of insulator proteins arrayed along an insulator may be less important for enhancer blocking activity. This difference can account for why DF and CDF have more enhancer blocking activity than does EF, whereas EF has more LR pairing activity (summarized in Figure 7). The GH region increases both LR pairing and enhancer blocking when added to EF, although these activities are still not as strong as those of DEF, with its Su(Hw) site intact (compare EF, EFGH, and DEF in Figures 2, S1, and 5a). Overall, while Su(Hw) provides a substantial part of *homie*’s pairing and enhancer blocking activities, other combinations of *homie*-bound insulator proteins apparently provide substantial activity as well.

Together, our data speak to the specificity of our LR pairing assay, which reflects the ability of *homie* to specifically pair with copies of itself when the two are located at considerable chromosomal distances from each other. The ability to pair with other insulators in the vicinity of the insertion site may be sufficient for enhancer blocking, but not for LR pairing. Alternatively, enhancer blocking may not require insulator pairing at all, even though it is clear that pairing can affect which interactions are blocked and which are facilitated by insulators.

### Enhancer blocking vs. PRE blocking

We compared the enhancer blocking and PRE blocking activities of an extensive set of *homie* sub-elements. These comparisons provide a clear indication of a mechanistic distinction between enhancer blocking and PRE blocking by *homie*. As Figure 5a shows, the D region has clearly more enhancer blocking activity than E, which has a small but significant amount, while the opposite is true in the PRE blocking assay, where E has strong activity, while D is considerably weaker (Figure 6). One possible explanation is that PRE blocking only requires the introduction of an extended nucleosome-depleted region in 1-dimensional space, along the chromosome, while enhancer blocking may require an additional ability to suppress looping in 3 dimensions between enhancers and promoters (reviewed in Bushey et al. 2008; Gaszner and Felsenfeld 2006).

### Su(Hw) and Cp190 motifs

Early studies found a genetic interaction between Su(Hw) and Cp190 (Pai et al. 2004), and suggested that the DNA binding insulator proteins BEAF, CTCF, and Su(Hw) dictate DNA sequence specificity, while Cp190 engages in protein-protein interactions through its BTB/POZ domain (Vogelmann et al. 2014). Although Cp190 has a zinc finger domain (Pai et al. 2004), it is unclear whether it can directly bind DNA (Vogelmann et al. 2014). Genome-wide analysis showed that both Su(Hw) and Cp190 are localized to *homie* and *nhomie* (Baxley et al. 2017; Cuartero et al. 2014; Negre et al. 2010; Schwartz et al. 2012; Soshnev et al. 2012; Wood et al. 2011). Indeed, we found DNA sequence motifs both for binding by Su(Hw) and “association with” Cp190 in *homie*’s D and E regions, respectively. Our data show that the Su(Hw) site has a major role in all three insulator activities, while the Cp190 site modestly affects LR pairing and enhancer blocking, but not PRE blocking activity (Figures 3, 4, S5, 5b, and 6b). A previous study using a mutation in the *Cp190* gene concluded that it affects enhancer blocking but not LR pairing by *homie* (Kaushal et al. 2022). It is possible that our LR pairing analysis is more sensitive than the one done in that study because of our extensive quantification and statistical analysis (Figure 3), but it is also possible that mutating the Cp190-associated site (which may be a Pita binding site, as described in Results) has effects beyond a reduced recruitment of Cp190. Conversely, other sites in *homie* could also contribute to Cp190 recruitment. Cp190 pull-down assays identified a number of physically associated insulator binding proteins (Kaushal et al. 2022), and the BTB/POZ domain of Cp190 is known to interact with several DNA binding insulator proteins, including Su(Hw), Pita, and CTCF (Golovnin et al. 2023). Other studies showed interactions between Cp190 and the DNA binding proteins Ibf1/2 (Cuartero et al. 2014) and M1BP (Bag et al. 2021). These studies suggest that Cp190 functions through interactions with other DNA binding proteins. All of the insulator proteins mentioned above localize to *homie*, based on genome-wide studies. The modest effects of mutating the Cp190-associated sequence in this study are consistent with other binding sites contributing to (indirect) Cp190 association with *homie*.

Previous studies showed that *gypsy*, containing 12 Su(Hw) binding sites (Parkhurst et al. 1988; Spana et al. 1988), is capable of self-pairing (Cai and Shen 2001; Muravyova et al. 2001), as well as having enhancer blocking and barrier activities (Geyer and Corces 1992; Holdridge and Dorsett 1991; Roseman et al. 1995; Roseman et al. 1993). As our data show, even though the Su(Hw) site in *homie* has a major role in each of its activities, *gypsy* does not show LR pairing with the *eve* locus. This distinction may arise from the strong requirement for pairing specificity in the LR pairing assay, as discussed above. How might Su(Hw) simultaneously harbor these different activities? A previous study showed that distinct zinc-fingers of Su(Hw) bind to different sequences in the core Su(Hw) site, and this could conceivably “activate” different functions, perhaps as a result of the recruitment of different cofactors (Baxley et al. 2017), or by facilitating interactions with a variety of other insulator-bound proteins. Consistent with this general notion, our data show that combining different sub-elements, such as adding F to DE (DEF vs. DE), adding 50 bp of the G region to D3EF (D3EFG5 vs. D3EF), or adding E to D (DE vs. D alone) increases both LR pairing and enhancer blocking activities (Figures 2, 3, 5a, and S1). Further analysis will be required to determine the specific interactions of these sub-elements with various insulator binding proteins, and how these interactions facilitate the three partially distinct insulator functions that we have studied here.

It has been suggested that the Su(Hw) consensus site should be extended beyond the core, because Su(Hw) contacts sequences flanking the core consensus binding site using parts of the protein that contribute differentially to different activities, including insulator activity (Baxley et al. 2017). Since the Su(Hw) site in *homie* is near the D-E junction, inverting EF separates one of the extended regions, an A/T-rich sequence, from the core site, which might cause a reduction in Su(Hw) binding or insulator activity, and thereby contribute to the observed loss of LR pairing. In the context used here, this A/T sequence in *homie* (TTTTT) is replaced by a sequence with G at one of the five positions (GATTA), both in the enhancer blocking and LR pairing assay vectors. The D region alone (which lacks the A/T sequence extension of the core Su(Hw) site) showed measurable enhancer blocking (Figure 5a) and PRE blocking (Figure 6b), and these activities are both totally dependent on the core Su(Hw) binding site (D vs. DΔSH), suggesting that the site has strong activity without the extended A/T sequence. Therefore, it seems unlikely that the difference in LR pairing between D3EF and D3FE is due solely to a change in the activity of the Su(Hw) site.

Several studies have shown that depletion of insulator proteins only moderately affects either boundary activities or TAD structure in cell lines, or during Drosophila embryogenesis (Chathoth et al. 2022; Gambetta and Furlong 2018; Kahn et al. 2023; Kaushal et al. 2022; Kaushal et al. 2021; Ramírez et al. 2018; Schwartz et al. 2012), including effects on the *eve* locus and surrounding genes (Van Bortle et al. 2012). For example, Cavalheiro *et al*. (2023) showed that removing or reducing individually the insulator proteins CTCF, BEAF, or Cp190 during *Drosophila* embryogenesis had no major impact on initial establishment of TAD structure, as assayed by Hi-C. This likely speaks to two issues. First, the plethora of insulator proteins in flies likely reflects a great deal of redundancy in their functions. Second, the effects of partially reducing insulator function are likely to be subtle, when viewed from a genome-scale perspective. For example, even when *nhomie* is completely removed from the *eve* locus, the changes seen at high resolution using Micro-C are noticeable, but not dramatic, as are the changes in gene function that result (Ke et al. 2024). Nonetheless, these changes are functionally important, and do seem to have been the object of a considerable amount of selection pressure, given that the genome is subdivided into thousands of TADs by insulator elements.

### Importance of the order of sub-elements

Our analysis of the D3EF combination uncovered another important feature of *homie* boundary activity.

We found that the order of the EF sub-elements in combination with D3 is important for LR pairing interactions, but not for enhancer blocking (Figures 3, S5, and 5b). Thus, while D3EF engages in LR pairing, inverting the EF sub-element to give D3FE completely disrupts LR pairing activity. This is consistent with a model in which the order of binding sites for chromosomal architectural proteins like Su(Hw), CTCF, and Pita plays a central role in determining the specificity and stability of boundary:boundary pairing interactions.

Many members of this particular class of polydactyl zinc finger proteins have self-interaction domains that can generate multimeric complexes, and potentially directly link boundary elements that have binding sites in common (Bonchuk et al. 2021; Fedotova et al. 2017). In this case, the ordering of those binding sites along the chromatin fiber would be expected to be important for both the specificity and stability of insulator pairing. The order of protein binding within the insulator is apparently less critical for its enhancer blocking function.

Inverting EF in D3EF to give D3FE could interfere with LR pairing at several levels. In order for the *eve* enhancers to activate reporter expression at a distance of almost 150 kb, across about a dozen intervening TADs, the transgene *homie* must recognize potential partners (*homie* and/or *nhomie*) in the *eve* TAD and “initiate” physical pairing (Bing et al. 2024). Changing the order of E and F may make it impossible for proteins associated with each sub-element to simultaneously interact with their partners in endogenous *homie* and *nhomie*, and the initiation step could fail. It is also possible that the initial interaction might involve only a subset of proteins associated with each boundary. In this case, factors associated with a single sub-element might be sufficient to initiate pairing, but steric hindrance arising from the mis-ordered sub-elements could prevent the formation of a sufficient number of physical links (a zippering up) between the boundary elements to generate a stable pairing interaction. Furthermore, live imaging experiments indicate that after pairing interactions are established and stabilized, activation of the transgene reporter by the *eve* enhancers requires a further “compaction” that appears to involve forming contacts between the *eve* enhancers and the promoter of the reporter (Chen et al. 2018). Though less likely, this step might also be perturbed by steric hindrance arising from misaligned sub-element-mediated contacts.

Several mechanisms have been proposed to explain how boundaries block enhancer-promoter interactions. Boundaries could act as roadblocks or sinks (Bi and Broach 1999; Blackwood and Kadonaga 1998; Blanton et al. 2003; Geyer 1997; Gohl et al. 2011), in which case blocking activity would be autonomous, and depend only on the functional properties of the proteins bound to D3EF and D3FE. Alternatively, insulation could be achieved by organizing the chromatin fiber into looped domains (Bing et al. 2024). In this case, the blocking activity of D3EF and D3FE would depend upon whether they can pair with boundaries in the neighborhood of the transgene. Clearly, a 1-dimensional (1-D) roadblock model for insulator function is insufficient, by itself, to explain the pattern of MicroC interactions within and between TADs (Bing et al. 2024), nor can it explain how E-P interactions between sequences that are 142 kb apart result in one reporter being activated, while the other, nearby reporter is not (Fujioka et al. 2016). More generally, a 1-D roadblock could explain enhancer blocking only to the extent that enhancers and promoters are communicating along the chromatin fiber, while we know that enhancers can loop to promoters without such “linear” communication. The sink model is not easily compatible with the directional activity of insulator-dependent enhancer blocking, since a sink should compete for enhancer-promoter interactions even when it is not located between them, acting more like a silencer than an insulator. So, it seems likely that enhancer blocking activity must be envisioned as a process in 3 dimensions that builds on the TAD organization of the chromosome, limiting the ability of enhancers and promoters to “find” each other by looping out the intervening DNA. Such a restriction on looping could be facilitated by the tendency of chromatin in different, insulator-defined TADs to coalesce together through copolymer co-segregation of similarly modified nucleosomes (Harris et al. 2023; Ke et al. 2024; Rowley and Corces 2018; Rowley et al. 2017).

A 1-D roadblock model, while it is not sufficient to fully account for enhancer blocking, might be sufficient for PRE blocking. Silent chromatin is thought to spread along a chromosome through mechanisms involving the recruitment of modifiers to existing silent domains, which then modify histones in nearby, mostly adjacent nucleosomes (Blackledge and Klose 2021). This difference could explain our results showing that PRE blocking is less demanding than enhancer blocking, in terms of its sequence requirements, in that smaller fragments of *homie* are sufficient for PRE blocking than for enhancer blocking (Fig. 7). On the other hand, PRE pairing can result in the spreading of repressive chromatin in *trans* (Kraft et al. 2022). So, even for PRE blocking, 3-D structures must be taken into account to fully understand the process. The requirements for LR pairing are more stringent still, which may reflect its requirement that distant sequences find each other and stably pair, in competition with closer pairwise arrangements of boundary elements, which would be more likely to find each other by random motion of the chromatin fiber.

## Materials and Methods

### Plasmid construction, transgenic fly production, and assay systems

Construction of the LR pairing assay vectors eZ (illustrated in Figure 1c; used in Figures 2, 5a, S1a-d, S3, and S8) and eZ-eG (illustrated in Figure 1d; used in Figures 3, 4, 5b, S1e, S2, S5, S6, S7, and S8b) were described previously (Fujioka et al. 2016; Fujioka et al. 2009). In short, the *lacZ* and *GFP* reporter genes are each driven by the *eve* promoter, and terminated by the *eve* 3’ UTR and polyA signal and the α-tubulin polyA signal, respectively. These vectors, carrying modified *homie* sequences, *gypsy*, or control λ DNA, were inserted into the attP site at –142kb relative to the *eve* locus (Fujioka et al. 2009). The -142 kb attP site is in the 5’ UTR of the *hebe* gene. Since the-142kb attP site was introduced into the chromosome using P-element insertion, both the 5’P-element end (5’P) and the 3’P-element end (3’P) are present (Figure 1c, d), and since these constructs were inserted into the attP site using recombinase-mediated cassette exchange (RMCE) (Bateman et al. 2006), they can be inserted in either orientation (Z5 or 3Z). Transgenes that carry wild-type *homie* interact with the *eve* locus and can express a reporter gene in a partial *eve* pattern, independent of their orientation, Z5 or 3Z (Bing et al. 2024; Fujioka et al. 2016; Fujioka et al. 2009). Modified *homie* sub-elements and combinations DE, DF, EF, D, DΔSH, E, EΔCp190 and F in the eZ vector carry 413 bp of λ phage DNA (λ DNA) as a spacer between the *eve*-*lacZ* promoter and the *homie* fragment.

The same constructs were used to analyze enhancer blocking activity. As diagrammed in Figure 1d, the *hebe* ventral midline enhancer is located upstream of the attP site in the chromosome. When the construct was inserted in the Z5 orientation, the test fragment is between the enhancer and *lacZ*. Wild-type *homie* prevents activation of *lacZ* by the ventral midline enhancer, while λ DNA is unable to do so, resulting in midline expression (Fujioka et al. 2016; Fujioka et al. 2009). On the other hand, in the 3Z insertion, the test fragment is not between the enhancer and *lacZ*, so the enhancer blocking activity of 3Z insertions cannot be determined with this assay (Figure 1c).

In order to localize the *hebe* midline enhancer, the 1^st^ intron of the *hebe*-RD transcript was split into three regions (Figure S8a: *hebe*-1, 2R:9829859 – 9832120; *hebe*-2, 2R:9831531 – 9834080; and *hebe*-3, 2R:9833624 – 9836462, based on genome assembly dm6 coordinates). To reduce the chance of missing an activity that spans the junction of two fragments, overlapping regions were tested. The regions were cloned into the eZ vector and inserted into the attP site at cytological location 74A2 (Fujioka et al. 2013). Enhancer activity was assessed as *lacZ* expression in a *hebe*-like pattern, using *in situ* hybridization.

The *eve* pseudo-locus construct used for analyzing PRE blocking activity was described previously (Fujioka et al. 2013). In short, the region located between either –6.4 kb (Figure S9) or –6.6 kb (Figure 6a) and +11.4 kb relative to the *eve* transcription start site was modified by replacing the region from +167 bp to +1.3 kb with the *lacZ* coding sequence and the *eve* poly-A signal. The 3’ end point of +11.4 kb, which is in the 3^rd^ exon of *TER94-RA* and -*RD*, was fused to the GFP coding region, followed by the *α-tubulin* poly-A signal. The attP sites used for this analysis (Fujioka et al. 2013) are at cytological locations 95E5 (Figure S9a) and 74A2 (Figure 6a and S9b). The smaller construct used to analyze PRE blocking activity (Figure 6b) is the same as the pseudo-locus, except that the sequence upstream of +8.4 kb was removed. The constructs using this vector were inserted into a MiMIC attP site, Mi{MIC}Drgx [MI04684] (Venken et al. 2011) (cytological location 24B1, used in Fujioka et al. 2021). As illustrated in Figure 1a, the *eve* 3’ PRE is located between +8.4 and +9.2 kb, just upstream of *homie* (Fujioka et al. 2008). In both constructs, the *homie* region from +9.2 to +9.8 kb (ABCDEF) was replaced with modified *homie* and 500 bp of λ DNA. When wild-type *homie* is in this position, it prevents the spreading of PRE-dependent repression activity into the *TER94-*driven *GFP* gene, and *TER94-GFP* is expressed strongly in ovaries (Fujioka et al. 2013). When λ DNA is in this position, it fails to prevent spreading of repression, and *TER94-GFP* is repressed. In order to maintain approximately normal positions relative to the *eve* PRE, modified *homie* derivatives DEF, CDE, CDF, and CEF each carry 217 bp of λ DNA, while AB, DE, DF, EF, C, D, E, F, DΔSH, and EΔCp each carry 413 bp of λ DNA, between the 3’ PRE and *homie*.

### Analysis of transgenic lines

*In situ* hybridization was performed based on previously published methods (Kosman et al. 2004), except that RNA was visualized using a histochemical reaction. Stage 4-7, stage 4-11, and stage 12-15 embryos were collected separately for several days. These samples were combined to make 60μl embryos per sample (so that stage 4-15 embryos are present in each sample). DIG-labeled antisense RNA probes against *lacZ* or *GFP* were produced using T7 RNA polymerase with DIG-RNA labeling mixture (Roche).

Antisense RNA was visualized using alkaline phosphatase-conjugated anti-DIG antibody (Roche), using CBIP and NBT as substrates (Roche). Once color was developed, which was determined by a positive control’s expression level (CDEF or DEF fragment in a transgene), the reactions of all samples were stopped simultaneously. Stained embryos were washed with ethanol to remove pinkish color, resulting in intensified dark blue color. Ethanol washing also prevents the bleeding out of pinkish color after embryos are mounted in Fluoromount (Southern Biotechnology). Each set of experiments was carried out with the positive control and experimental samples in parallel to minimize experimental variation. Images were obtained using a Zeiss Axioplan2 microscope with the same camera settings each time. Each experiment was performed at least twice, with independent *in situ* procedures. Representative expression is shown in the figures.

smFISH (Little and Gregor 2018; Trcek et al. 2017) was performed as previously described (Ke et al. 2024). Dechorionated embryos (stages 14-16) were fixed in 5mL 4% paraformaldehyde in 1X PBS and 5mL heptane for 15min with horizontal shaking. After devitellinization, embryos were washed 2X with 1mL of methanol. Methanol was then removed and replaced by PTw (1X PBS with 0.1% Tween-20) through serial dilutions of 7:3, 1:1, and 3:7 methanol:PTw. The embryos were then washed 2X in 1mL of PTw and 2X in 1mL smFISH wash buffer (4X SSC, 35% formamide, and 0.1% Tween-20), and incubated with ∼5nM coupled smFISH probes (Biosearch^TM^) in hybridization buffer (0.1g/mL dextran sulfate, 0.1mg/mL salmon sperm ssDNA, 2mM ribonucleoside vanadyl complex, 20μg/mL RNase-free BSA, 4X SSC, 1% Tween-20, and 35% formamide) for 16h. Embryos were then washed 2X for 2h in 1mL smFISH wash buffer, followed by 4X 30min washing in 1mL PTw. For DAPI/Hoechst staining, the embryos were stained with 1ug/mL DAPI or Hoechst in PTw for 15min, then washed 3X for 5min with 1mL PTw. Finally, the embryos were mounted on microscope slides with Aqua PolyMount and a #1.5 coverslip for imaging.

For assaying LR pairing activity, we assessed the number of cells expressing *lacZ* in an *eve* pattern, and their intensities, at stages 5–8, when the reporter is expressed in stripes, and when *eve* is tissue-specifically expressed (mesoderm, APR, and CNS) at stages 11–13. First, we made an overall assessment of the activity ranking of the reporter based on at least 2 independent experiments. When *lacZ* expression levels and cell numbers were similar between reporters, stage 5 embryos from sets of embryos stained in parallel were subjected to cell counting and analysis. Transgenes with expression similar to that of λ DNA (*i.e*., ABCD, CDF, DF, D, E, F, DEF-ΔSH, D3FE, and *gypsy*) were excluded from this analysis, as they had few, if any, cells expressing the *lacZ* reporter in an *eve*-like pattern. For those chosen for detailed analysis (see Figures 2 and 3), we manually counted *lacZ*-expressing cells, and measured the staining intensity of each of those cells using the ROI tools of Image J software (Schneider et al. 2012). When a cell’s shape was recognizable from the *lacZ* pattern, they were counted as expressing. To obtain the average intensity of expression per expressing cell for an embryo, we first chose 10 cell-sized locations each, outside the embryo and inside (where there were no *lacZ*-expressing cells), and measured the “background” light intensity. The outside background was taken as the maximum light intensity, representing the lowest possible expression signal. The average light intensity of each expressing cell within each embryo, as well as the average of the “inside” background, were each subtracted from this maximum light intensity to get the signal strength for each cell and the average signal background, respectively. The average signal background was then subtracted from the signal strength to get the signal intensity above background for each cell, and these individual cell signals were averaged to get the average signal intensity above background per expressing cell for each embryo. For this cell count and intensity analysis, we took the following precautions. First, the expression of *eve*, and therefore of the *lacZ* reporter, is rapidly changing around stage 5, so small differences in developmental timing can affect the results. So, for each sample, we used Nomarski microscopy to identify closely matched embryos based on the extent of invagination of cell membranes as cellularization of the blastoderm proceeds. Also, the viewing angle of mounted embryos can affect the number of visible cells. We compared one lateral side of each embryo to the other side (two focal planes), and chose embryos in which the cell numbers were similar in the two focal planes. The clearer of these two focal planes (usually the closer one) was used for the counting. After applying these restrictions, 5-7 embryos/construct/staining could be analyzed. In order to increase the statistical power of the analysis, data from 2-3 independent stainings were combined. For this purpose, and to standardize the data to provide for easier comparison of activities in the different assays, values for each set of constructs were linearly scaled relative to those of CDEF (positive control = 100) and λ DNA (negative control = 0). Sample sizes are given in Figure S11 (including total number of cells counted and quantified, total number of embryos analyzed, and number of different independent experiments/stainings included in the analysis, for each construct), along with the results of pairwise t-tests of the differences in LR pairing for all constructs quantified, for both number of cells expressing per embryo and average intensity of expression per expressing cell per embryo.

For analyzing enhancer blocking activity, late stage 12 and 13 embryos were separately subjected to counting of CNS ventral midline cell clusters expressing the reporter gene. When cell shapes were visible, it was counted as an expressing cluster. Embryos from at least 2 independently stained samples were analyzed, and the results were combined. The distribution of expressing cell clusters per embryo was graphed as a box-and-whiskers plot, and the pair-wise significance of differences (p-values) between constructs was calculated using the t-test function in Excel (Microsoft). Expression in the CNS for each line is shown in Figure S8b. The number of embryos analyzed for each construct and stage is given in Figure S11.

For analyzing PRE blocking activity, ovaries were dissected from adult female flies aged 18–24h at room temperature. Live GFP images of ovaries were obtained using the same camera settings throughout, except as noted in the figures. In order to show consistency of GFP expression among ovarioles, several ovarioles are shown in each picture. In order to quantify *GFP* reporter RNA expression, ovaries were subjected to RT-PCR (Figures 6 and S9). Total RNA was extracted from 5-8 pairs of ovaries dissected from 18–22h adult females using an RNA extraction kit (Roche). 50ng of total RNA was used to make cDNA using the Transcriptor First Strand cDNA Synthesis kit with random primer (Roche). One μl out of 50μl cDNA solution was used for each quantitative PCR (qPCR) reaction, each time performed in triplicate. Data are shown as *GFP* expression normalized to endogenous *TER94* RNA expression, similarly quantified in each sample. Each set of lines was analyzed in 3 independent assays, and their averages and standard deviations were graphed. The pair-wise significance of the difference (p-value) between these construct averages was calculated using the t-test function in Excel (Microsoft). Primers used were, for *GFP*: GGGCACAAGCTGGAGTACAACTACAA and TGGCGGATCTTGAAGTTCACCTTG, and for *TER94*: TGAAGCCACCGCGTGGTATTCTTA and TTTGGACATGATCTCCGGTCCGTT.

Statistical analysis (t-test results) for these three assays is given in Figure S11.

### Binding site analysis

Binding site searches was done using MacVector software (MacVector Inc). We used DNA binding motif logos presented in genome-wide studies. When the frequency of nucleotide occurrence was similar at a position, it was represented by the appropriate single letter code. In order to identify motifs in *homie*, several mismatches were allowed in the search. Su(Hw) DNA binding motifs used were YWGCATACTTTT (Negre et al. 2010) and NWWWWNYRTWGCATACTTTTNKGSDB (Baxley et al. 2017). The Cp190-associated consensus sequence used was GGTTBDWRWMYYNGCTD (Cuartero et al. 2014). Mutations were introduced at base pairs that appear at high frequency in the motif. Additionally, a sequence motif for Pita binding, TAGCVDRKDHNHVMWCC (Maksimenko et al. 2015) was searched for. The results are shown in Figure S10. For cloning purposes, BamHI and HindIII restriction enzyme recognition sequences were added for the Su(Hw) and Cp190 site mutations, respectively. The ChIP-seq data shown in Figure S4 for Su(Hw) (Wood et al. 2011), Cp190 (Wood et al. 2011), and Pita (Zolotarev et al. 2016) were visualized using IGV (Thorvaldsdóttir et al. 2013).

## Data Availability

All data underlying this publication are included in the text and figures.

## Acknowledgements

This work was supported by NIH grants to Paul Schedl (5R35GM126975) and James B. Jaynes (1R01GM137062). We thank Qing Liu for excellent technical assistance.

## Supplemental Figures

**Figure S1.**
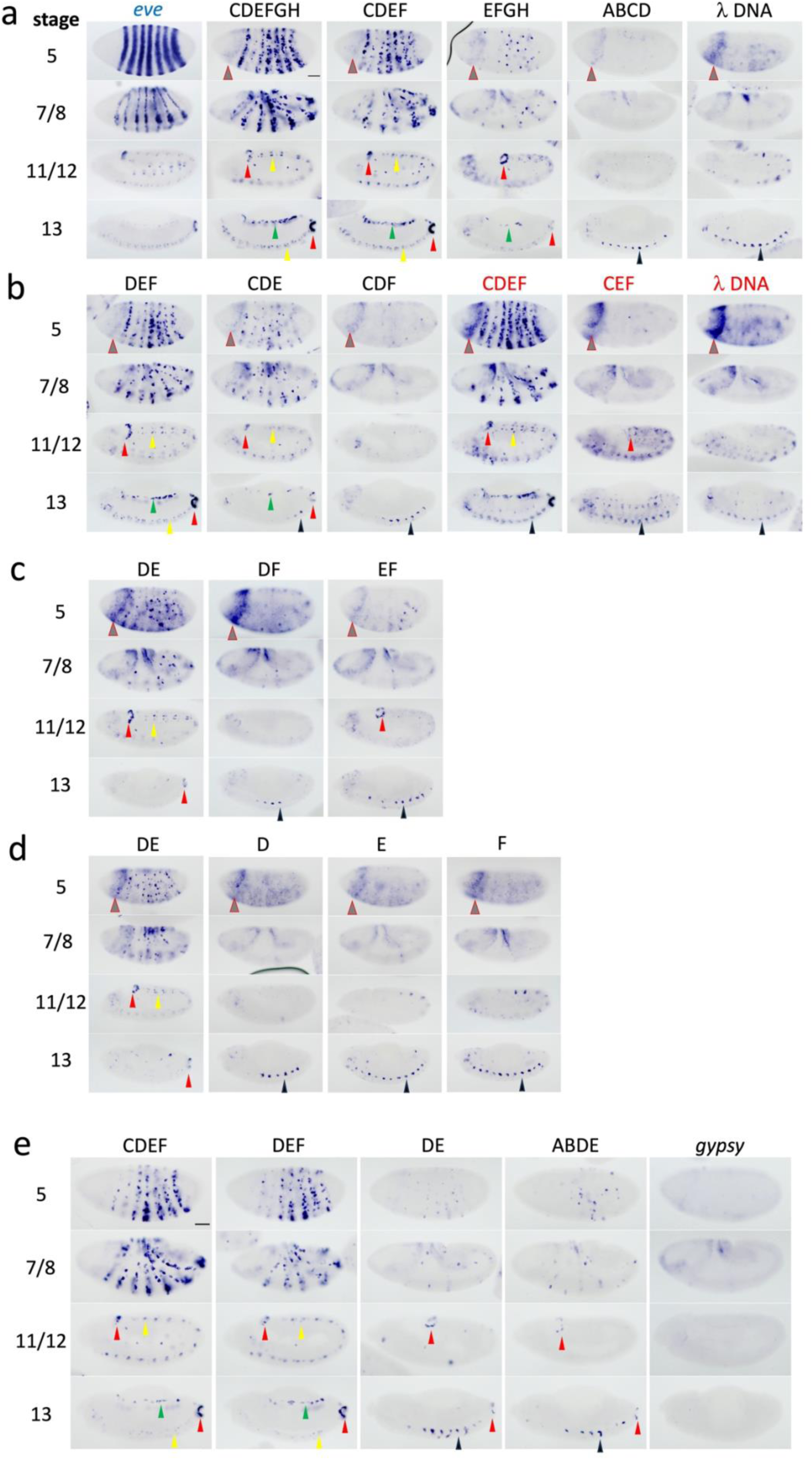
300 bp of *homie* (DEF) contains most of its long-range pairing activity. The previously described 800 bp “full-length” *homie* (Fujioka et al. 2009) was further dissected, by dividing it into roughly 100 bp segments (A–H, see Figure 1b) and testing them in our long-range (LR) interaction assay. For reference, expression of endogenous *eve* RNA is shown at the left in **a** (“*eve*”). For each tested region (see map in Figure 1b), *lacZ* reporter expression from eZ-vector transgenes (**a-d**) or from eZ-eG-vector transgenes (**e**) at embryonic stages 5, 7/8, 11/12, and 13 is shown. The color of the name of the *homie* derivative at the top indicates the transgene orientation in the chromosome: black is Z5, red is 3Z (see maps in Figure 1c, d). **λ DNA**: 500 bp of phage λ DNA, which serves as a negative control. ***gypsy***: a 349 bp *gypsy* transposable element-derived sequence was tested in the eZ-eG transgene in **e**. Red arrowheads: anal plate ring expression (APR); green arrowhead: mesodermal expression; yellow arrowhead: CNS expression; black arrowheads: *hebe*-like midline expression; and grey arrowhead with red outline: background expression seen most prominently in the 3Z orientation, and in the absence of a strong insulator (see text). *In situ* hybridizations shown in **d** and **e** were performed on a different day than those shown in **a–c**. Scale bar = 50μm.

**Figure S2.**
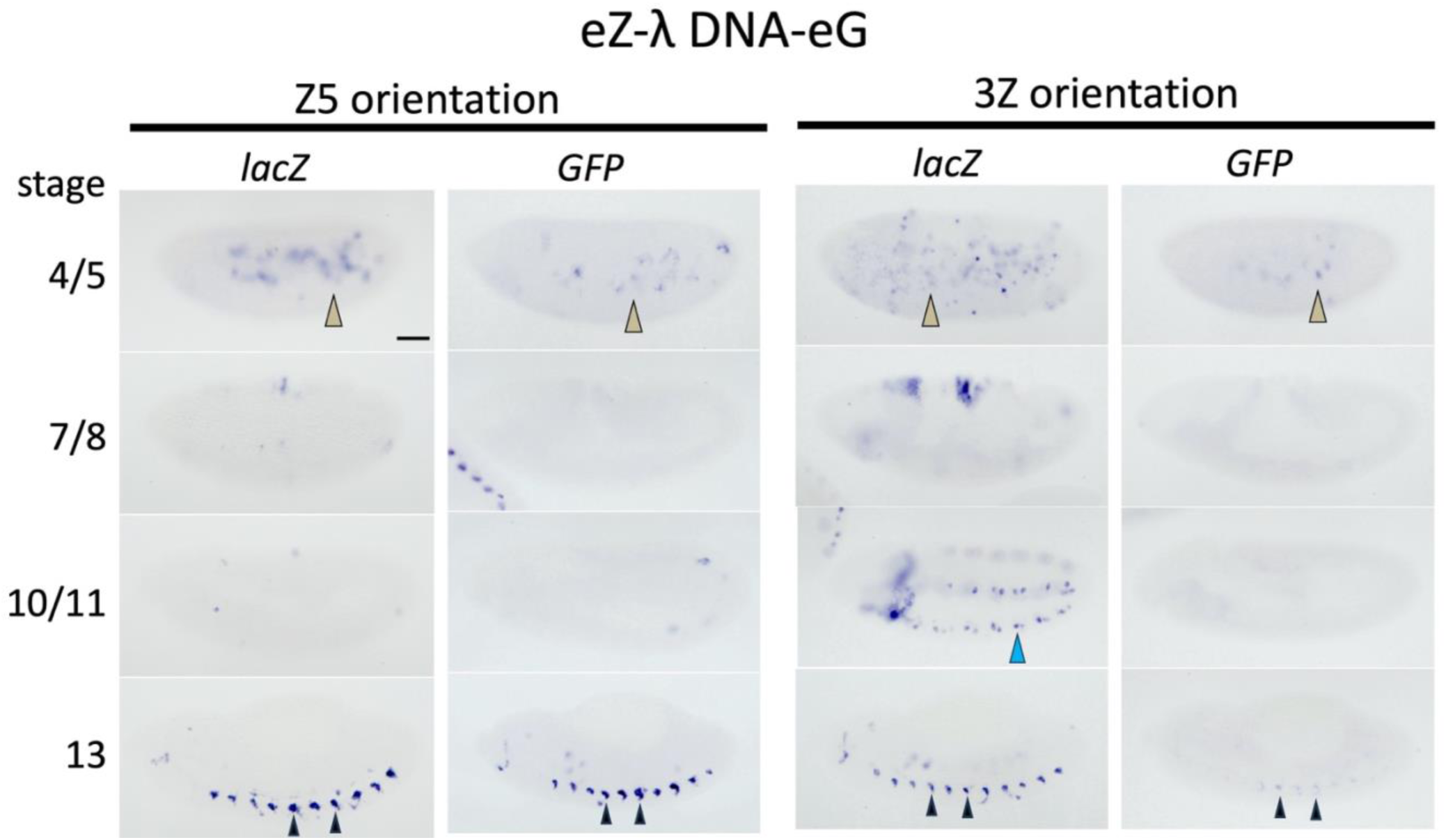
Negative controls for eZ-eG vector transgenes carrying *homie* derivatives, inserted at –142 kb relative to endogenous *eve*. Reporter gene expression (*lacZ* or *GFP* RNA, as indicated) from transgenes inserted in the indicated orientation (Z5 or 3Z), carrying 500 bp of λ DNA between the reporter genes. Embryos at stages 4/5, 7/8, 10/11, and 13 are shown, as indicated on the left. Non-*eve*-related expression is as follows: olive arrowheads outlined in black: expression from yolk nuclei; blue arrowhead: lateral expression; black arrowheads: ventral midline (*hebe*-like) expression. Scale bar = 50μm.

**Figure S3.**
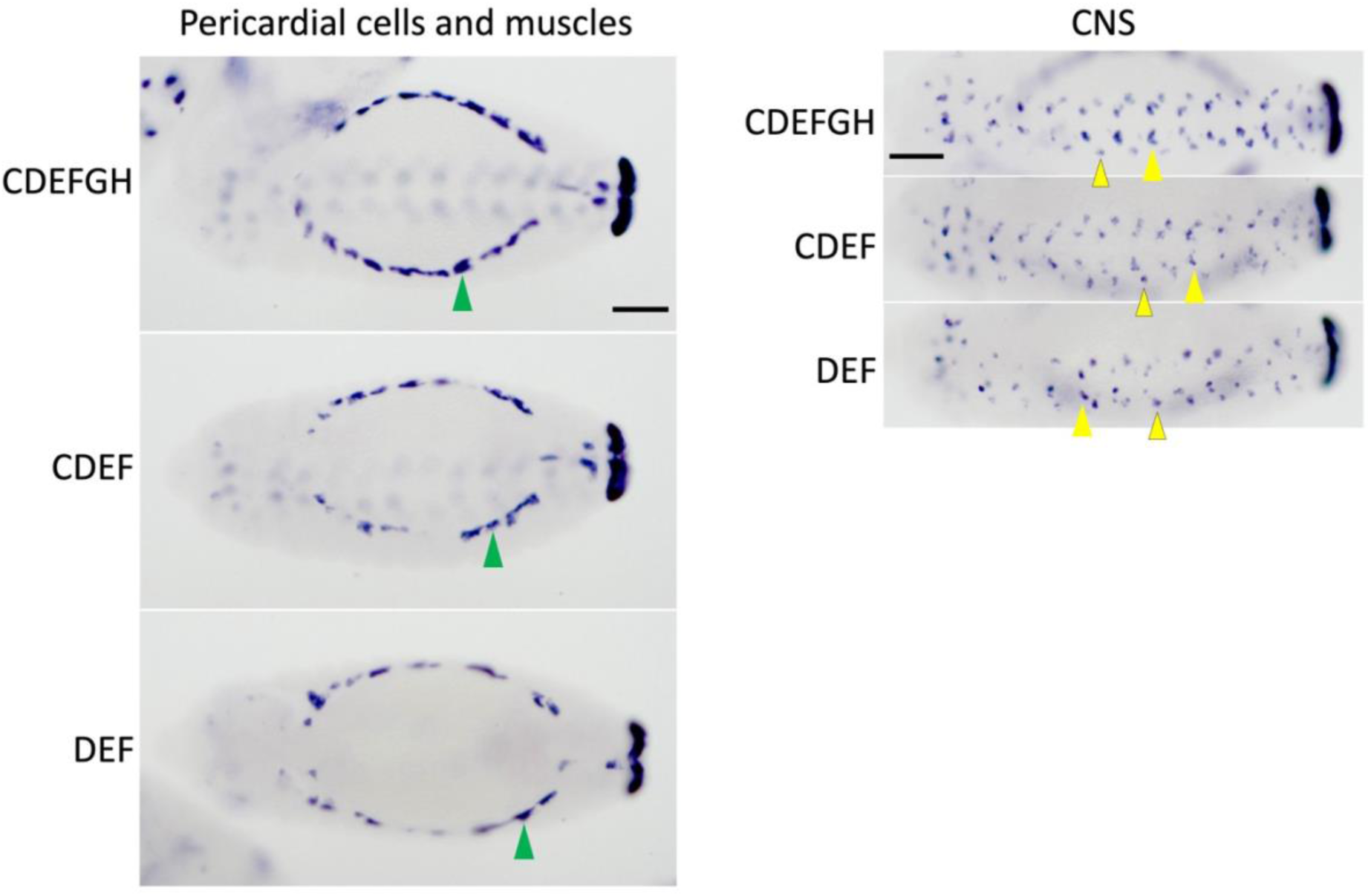
Expression of *eZ-lacZ* transgenes in *eve*-expressing embryonic mesoderm and CNS. *lacZ* RNA expression in transgenic embryos carrying the indicated *homie* derivatives are shown. Left panel: dorsal view of stage 13 embryos. Green arrowheads indicate mesodermal cells. Right panel: ventral view of stage 13 embryos. Yellow arrowheads indicate *lacZ* expression in CQ and EL (black outlined) neuronal precursor cells. Scale bars = 50μm.

**Figure S4.**
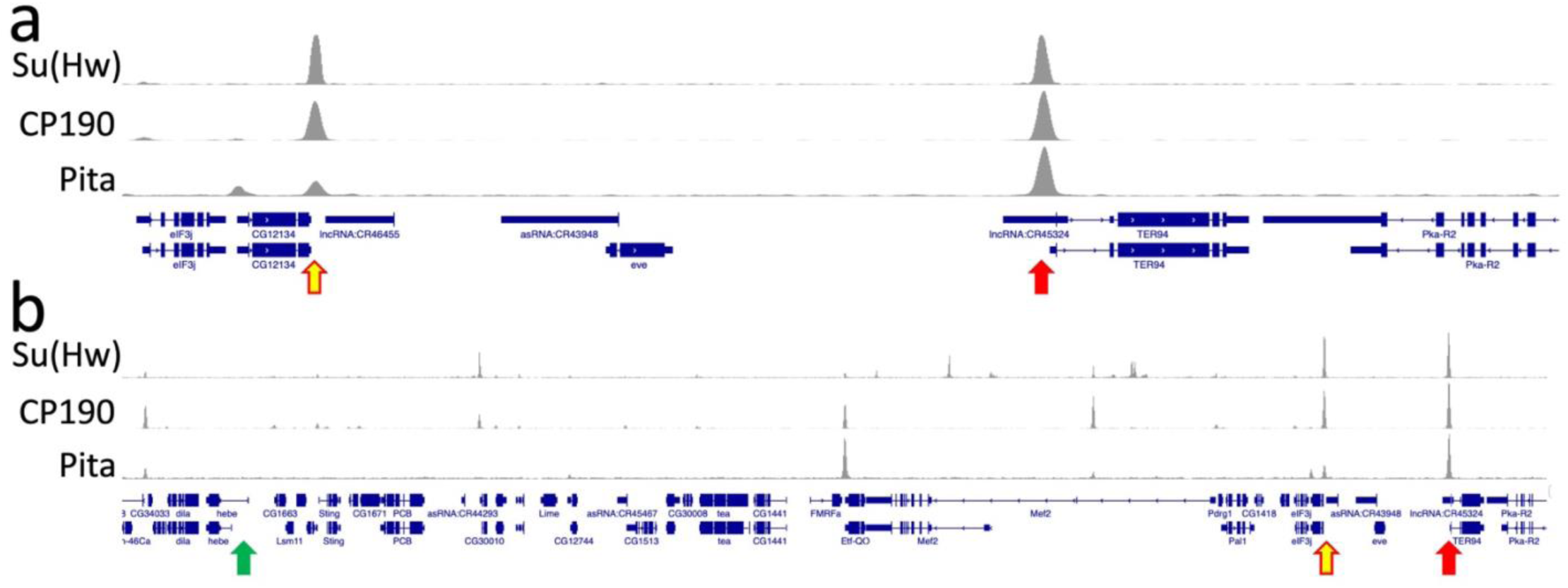
Su(Hw), Cp190, and Pita bind to *homie* and *nhomie*. ChIP-seq data from Wood *et al*. (2011) for Su(Hw) and Cp190, and Zolotarev *et al*. (2016) for Pita are shown. Yellow arrow with red outline: *nhomie*; red arrow: *homie*; green arrow: *hebe*.

**Figure S5.**
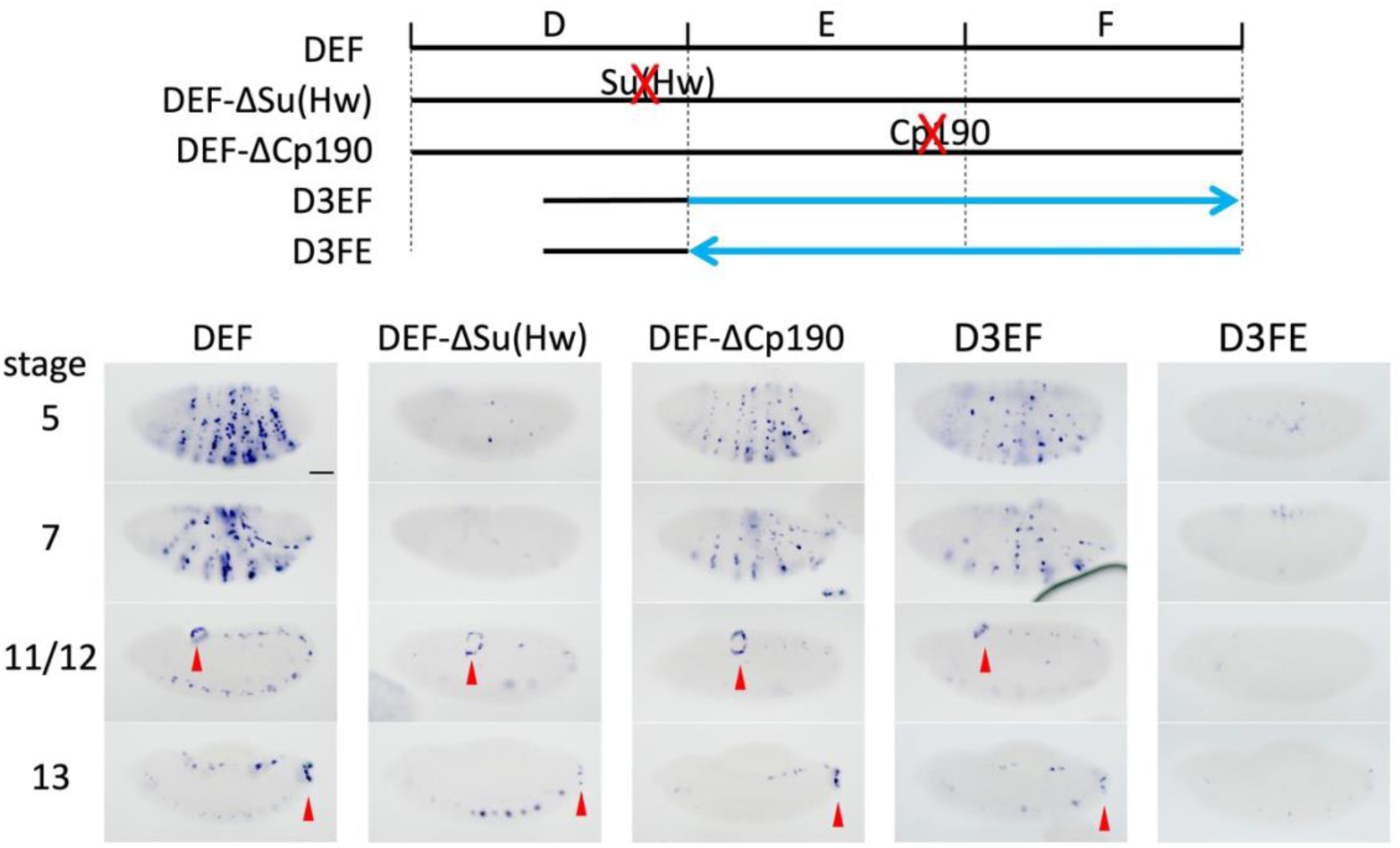
A consensus binding site for Su(Hw) and the order of sub-elements contribute to long-range pairing by *homie*. *lacZ* expression is shown from eZ-eG transgenes in the Z5 orientation at embryonic stages 5, 7/8, 11/12, and 13. The positions of Su(Hw) and Cp190-associated consensus sites are shown in the diagram at the top. “ΔSu(Hw)” and “ΔCp190” indicate versions with the site point-mutated (mutant sequences are given in Figure S10). In D3EF, the 5’ (left) half of D was removed from DEF (the Su(Hw) site remains intact), while in D3FE, the orientation of section EF was additionally reversed. The orientation of EF is indicated by the blue arrows in the diagram. Scale bar = 50μm.

**Figure S6.**
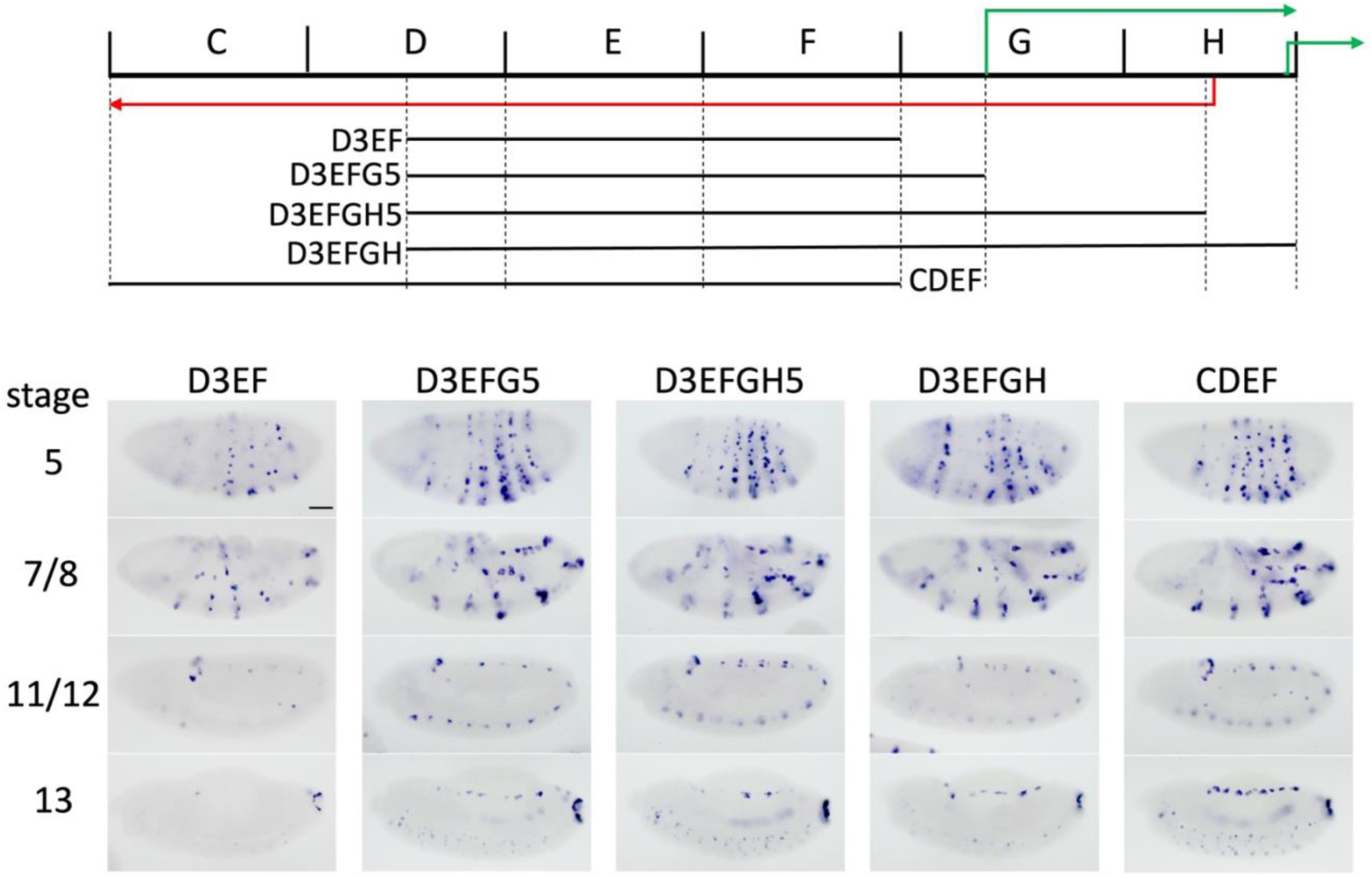
A sequence downstream of *homie* F contributes to LR pairing. **Top:** A diagram of the CDEFGH region. The green and red arrows show the directions of transcription of *TER94* and non-coding *CR45324* RNAs, respectively. Shown is *lacZ* expression from eZ-eG transgenes in the Z5 orientation at embryonic stages 5, 7/8, 11/12, and 13. The regions used for the analysis are shown in the diagram at the top. In D3EFG5, D3EF is extended to just downstream of the *TER94-RA* start site; in D3EFGH5, it is extended to just downstream of the *CR45324* start site; in D3EFGH, it is extended through the end of *homie* subregion H, which includes the *TER94-RD* start site. Scale bar = 50μm.

**Figure S7.**
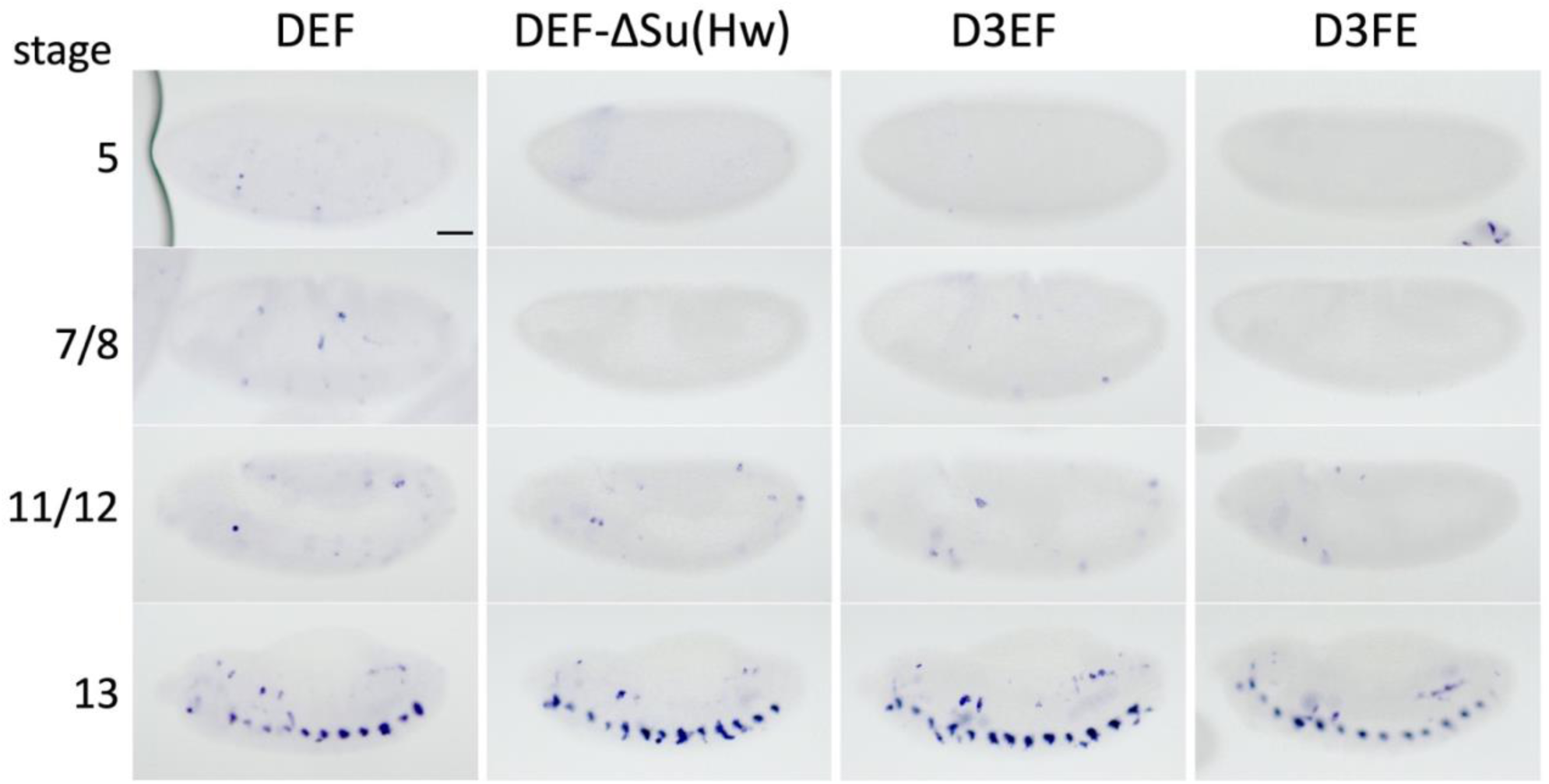
*GFP* RNA expression from transgenes used in. Figure 3a. *GFP* RNA expression in transgenic embryos carrying the indicated *homie* fragments in the eZ-eG vector in the Z5 orientation. Embryonic stages 5, 7/8, 11/12, and 13 are shown, as indicated at left. Scale bar = 50μm. Note that there is no *GFP* expression in an *eve* pattern, except for a few cells in DEF at stages 5 and 7/8 that are likely within endogenous *eve* stripes.

**Figure S8.**
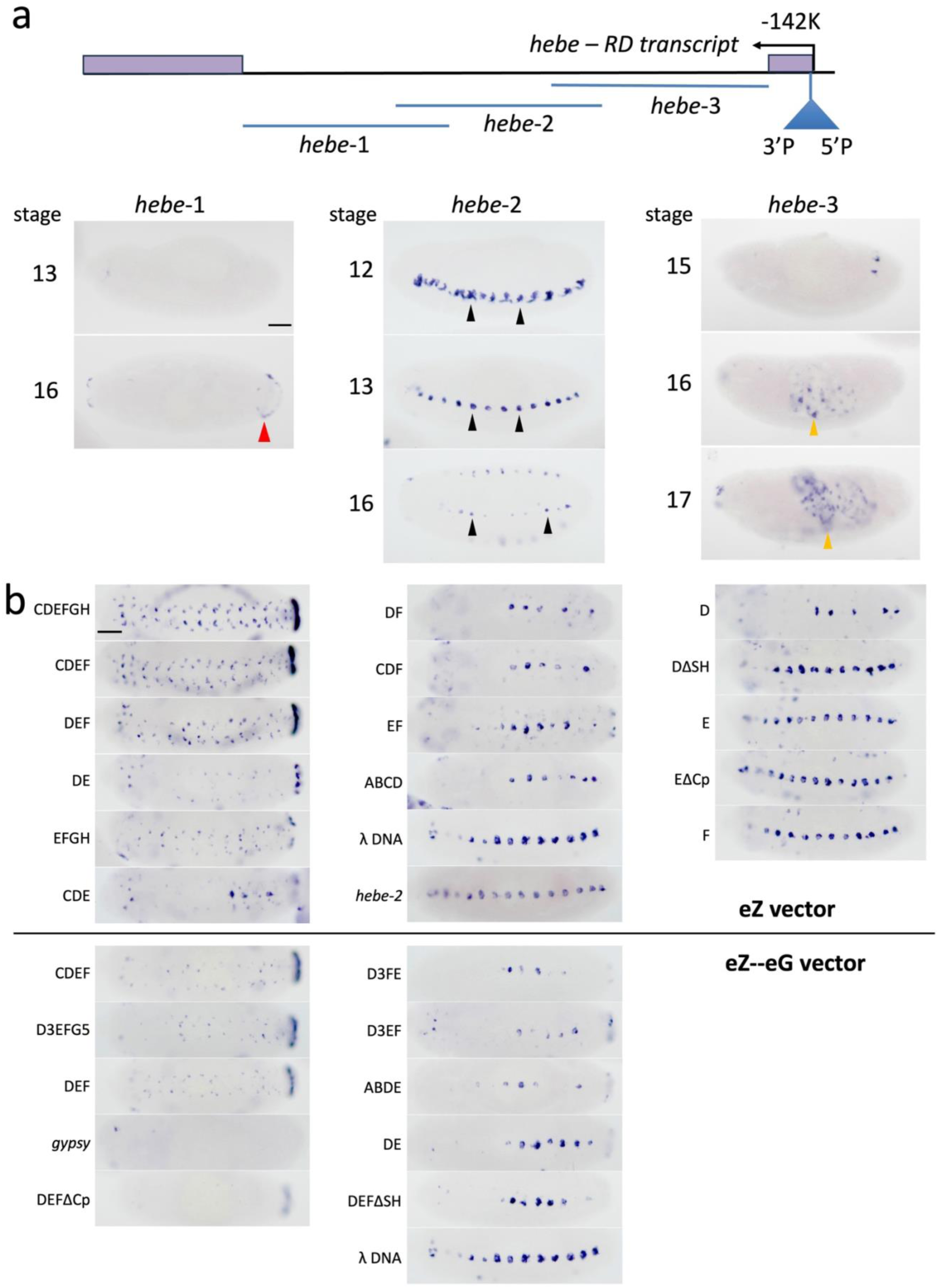
Mapping of *hebe* enhancers located near the transgene insertion site at –142 kb from endogenous *eve*, and ventral views of embryos carrying *homie* and derivatives. Scale bars = 50μm. **a.** The 1^st^ intron of *hebe* was analyzed for enhancer activity. In order to identify *hebe* enhancers, eZ-vector transgenes carrying DNA fragments *hebe*-1 (2.3 kb), *hebe*-2 (2.5 kb), and *hebe*-3 (2.8 kb) were inserted at the 74A2 attP site (on chromosome 3, where there is no interaction with the *eve* locus, which is on chromosome 2). The region analyzed is diagrammed at the top. *lacZ* expression driven by each fragment is shown below. The number to the left of each image indicates the embryonic stage. Red arrowhead: APR expression driven by *hebe*-1, which is seen only at stage 16 and later. Black arrowheads: ventral midline expression, strong at stages 12-13, fading by stage 16. Orange arrowheads: expression in the gut, seen at stage 16 and later. **b.** Ventral views of stage 13 embryos carrying the *homie* elements used in Figure 5. that in the “wt” pseudo-locus.

**Figure S9.**
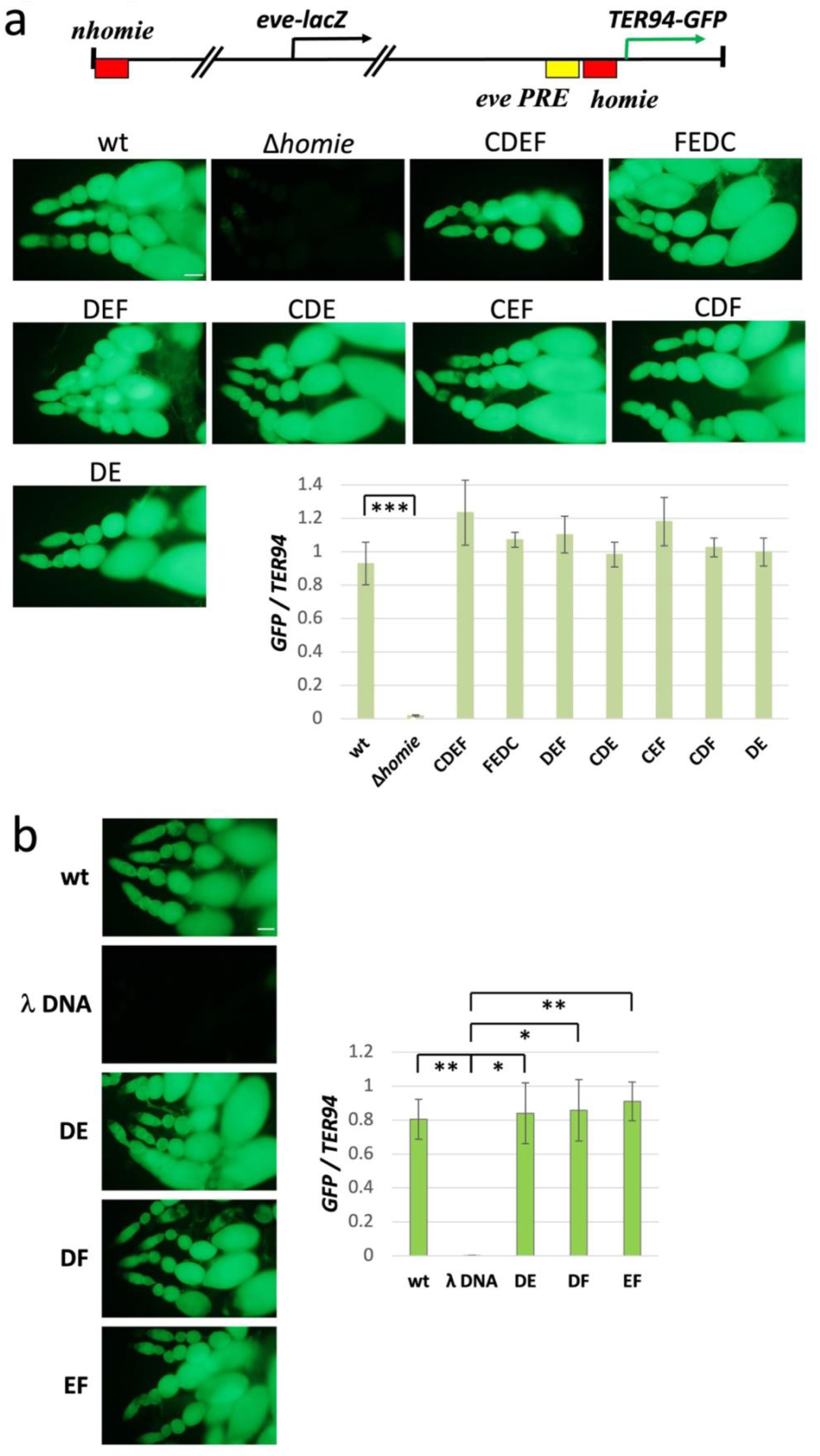
PRE blocking activities of *homie* sub- element combinations. Top: Diagram of the *eve* pseudo- locus transgene, inserted at either 95E5 (**a**) or 74A2 (**b**) (Fujioka et al. 2013). **Images:** Live images of *GFP* expression in dissected ovarioles. Scale bars = 50μm. **wt**: intact pseudo-locus. **Graphs:** quantification of *GFP* expression by RT-PCR. *GFP* RNA levels were normalized to endogenous *TER94* RNA. The averages and standard deviations of 3 independent data sets are shown. The results of pairwise statistical comparisons (t-tests as implemented by Microsoft Excel) are shown as brackets connecting each pair. Significance of the difference is indicated by the number of asterisks within each bracket: * = p < .05, ** = p < .01, *** = p < .001. **a. Δ*homie***: ABCDEF *homie* deleted; **CDEF, FEDC, DEF, CDE, CEF, CDF, DE**: the ABCDEF region of *homie* was replaced with each of these *homie* derivatives. Differences among *homie*-carrying transgenes were insignificant (p > 0.05 for each against wt). **b. λ DNA**: the same sequence used in Figure 2 replaced ABCDEF. **DE, DF, EF**: the ABCDEF region of *homie* was replaced with each of these *homie* derivatives. **a** and **b**: In each case, λ DNA was used to make the spacing between the 3’ PRE and the *homie* derivative similar to

**Figure S10.**
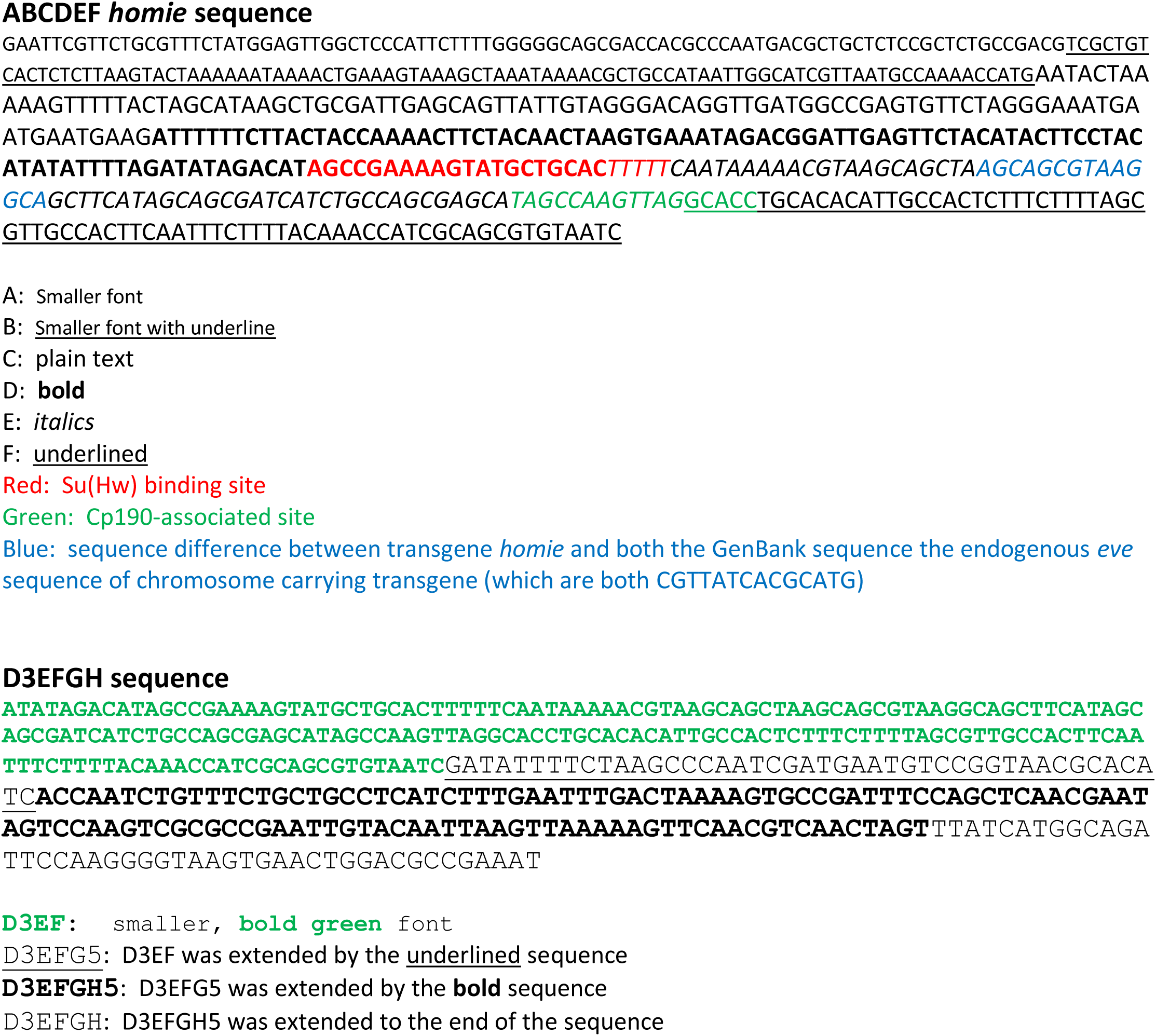
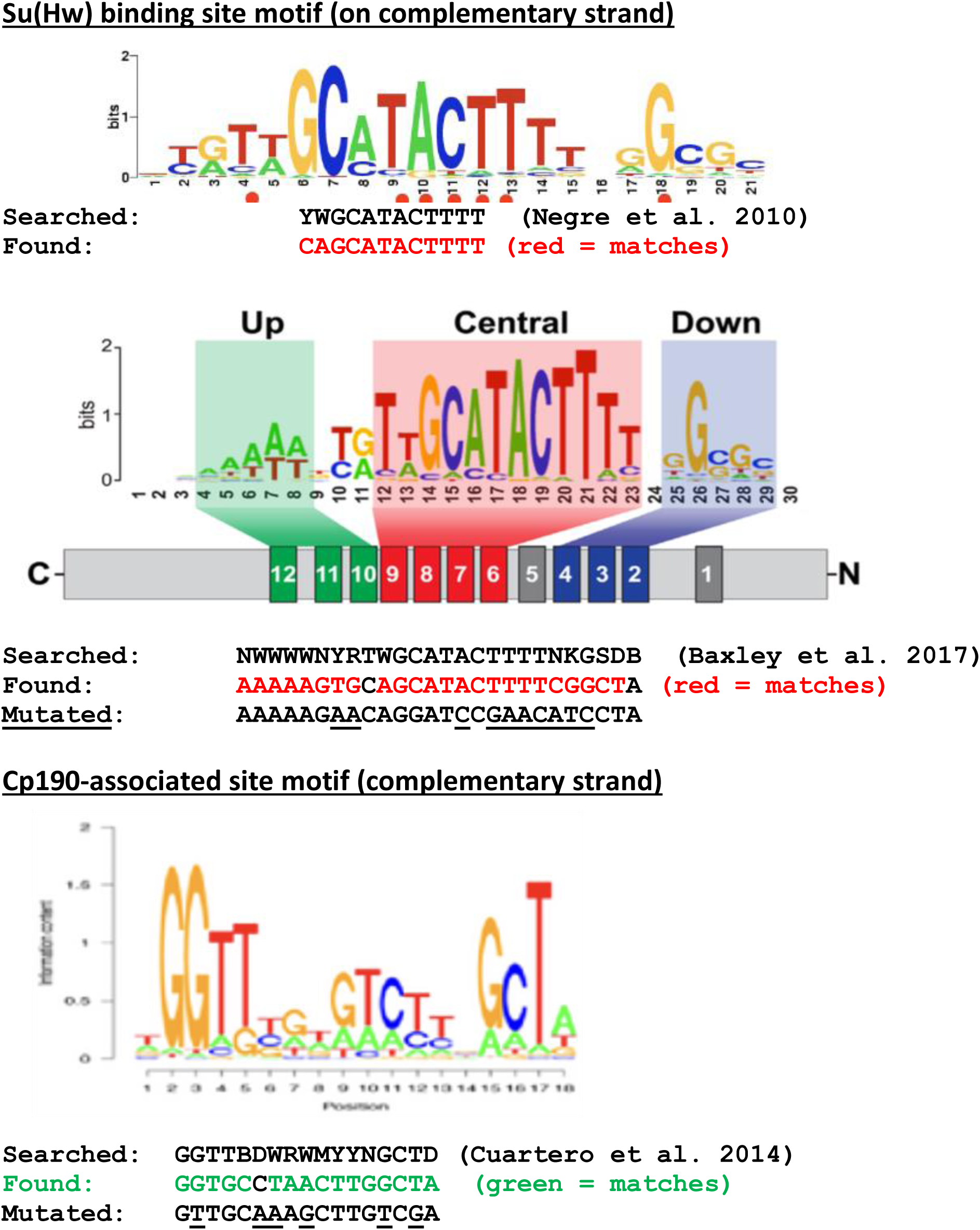
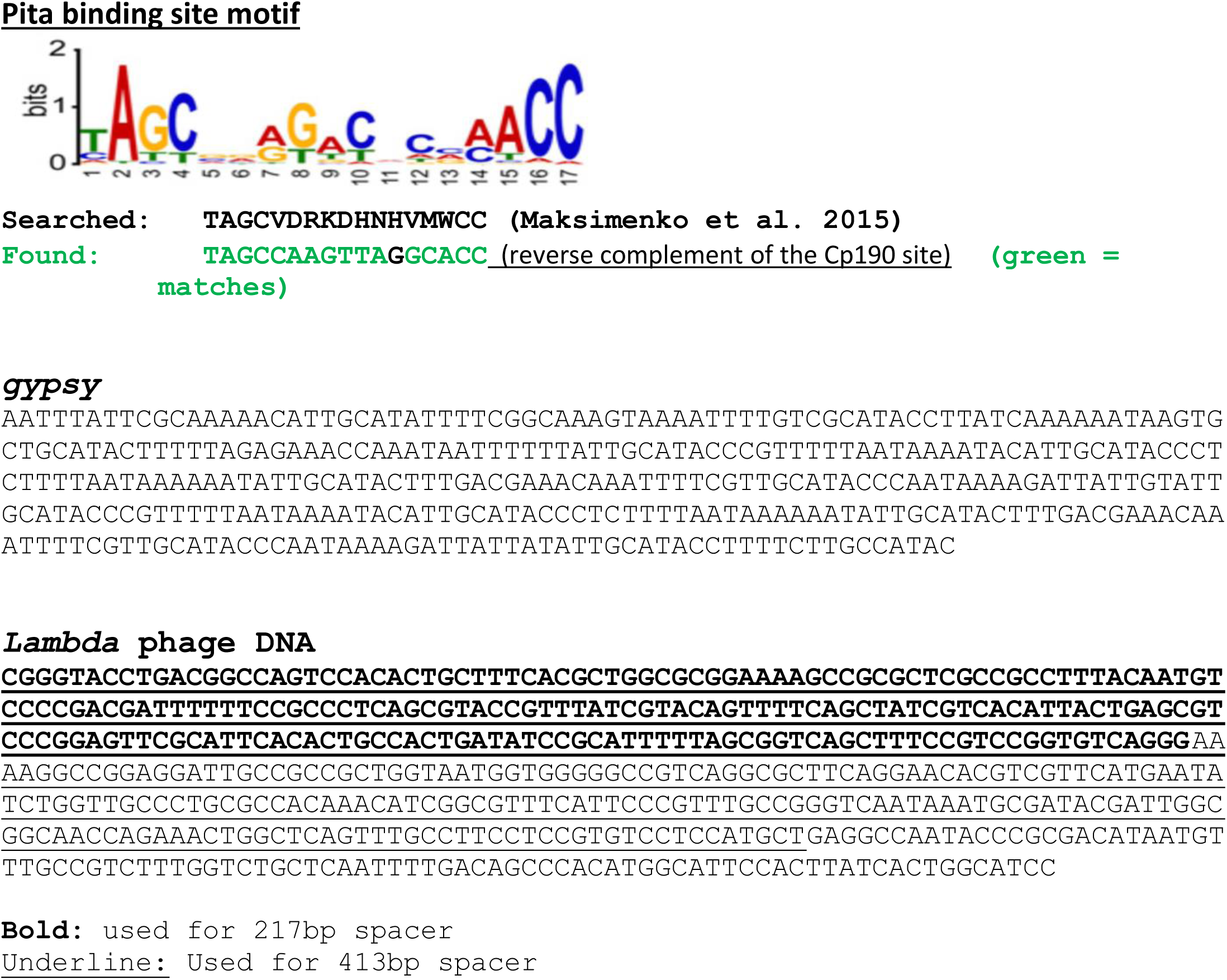
DNA sequences (*homie*, *gypsy*, and λ DNA) used in this study, and DNA sequence motif search sequences and results. See the main text and figure legends for how these sequences and search results were used.

**Figure S11.**
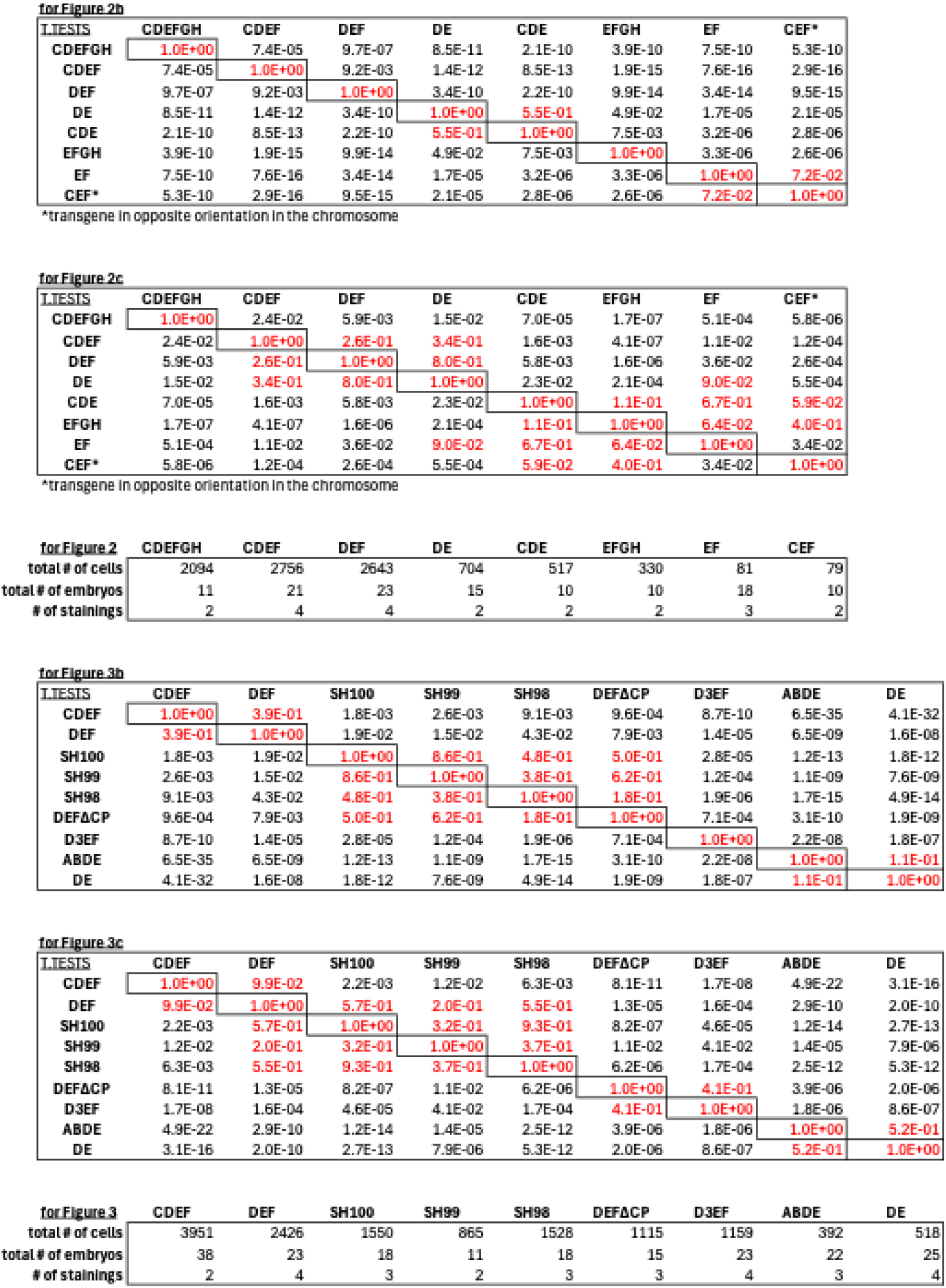
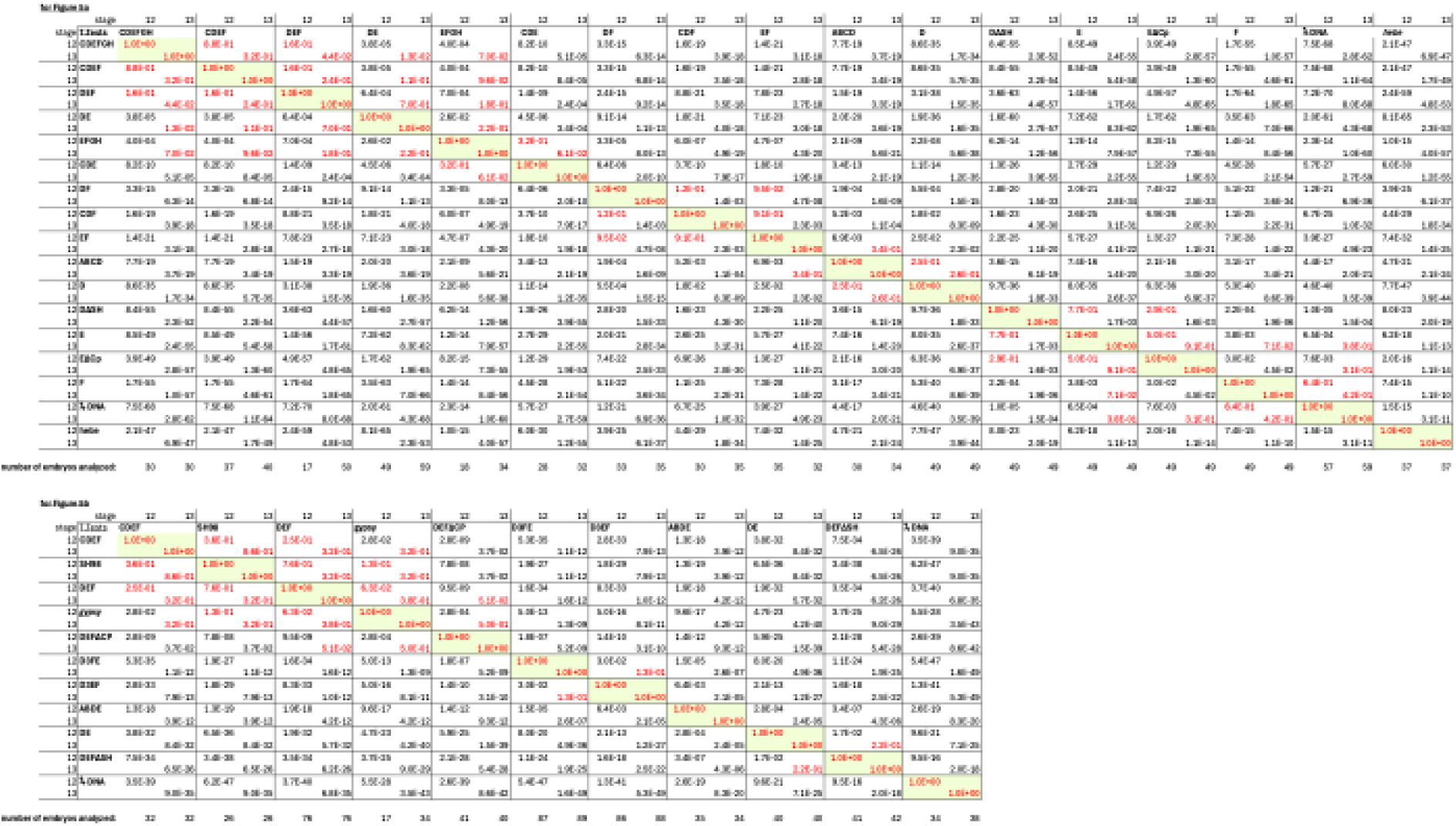
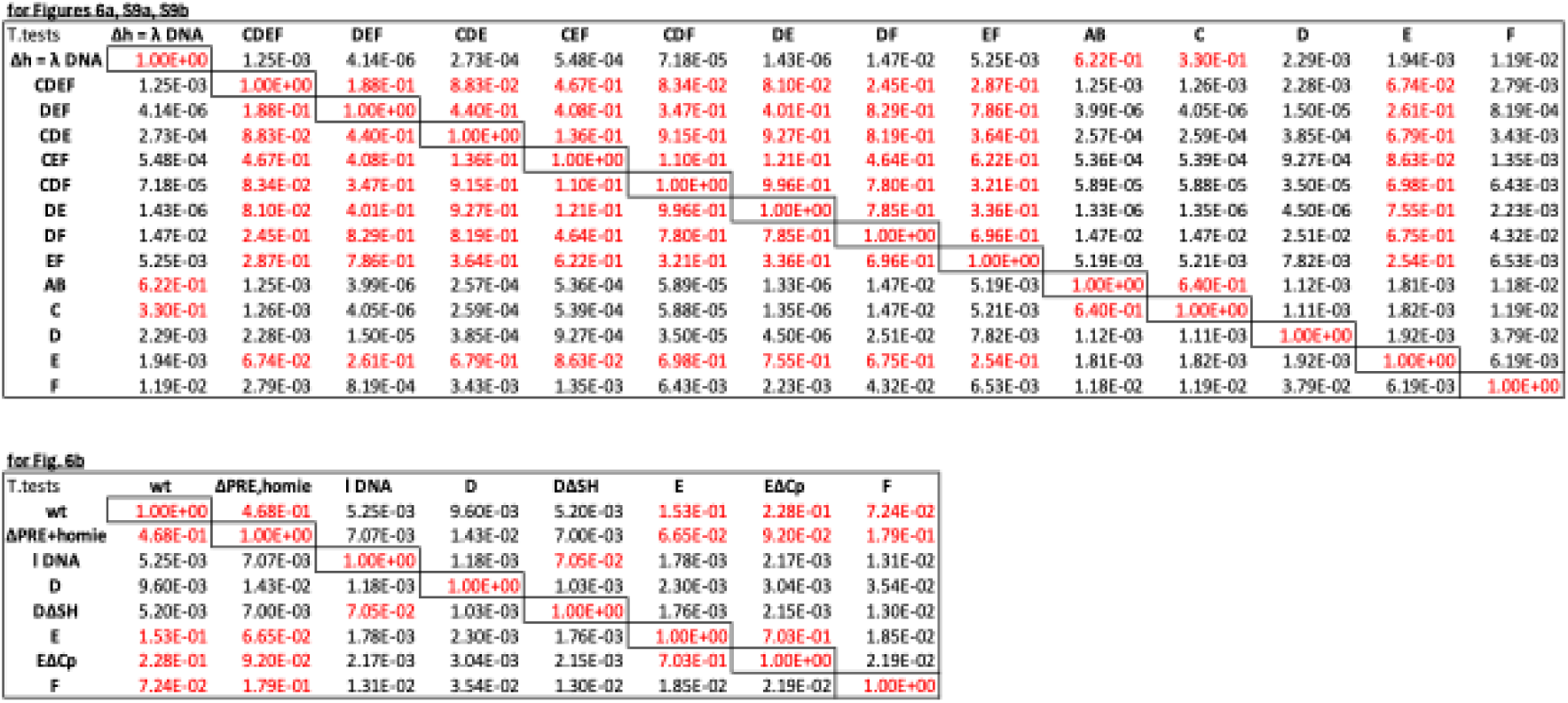
T-test results related to graphs in other figures, and sizes of LR pairing and enhancer blocking data sets. T-test grids show significance (p) values for pairwise comparisons of the indicated data sets (column label to row label: two-tailed t-test with unequal variances). Values in red represent no significant difference (*i.e.*, p > 0.05).

## Notes

### Competing Interest Statement

The authors have declared no competing interest.

### Summary of Updates

Additional data have been added that reinforce the main conclusions, and figures have been reorganized to present the data more effectively.

